# Global regulatory transitions at core promoters demarcate the mammalian germline cycle

**DOI:** 10.1101/2020.10.30.361865

**Authors:** Nevena Cvetesic, Malgorzata Borkowska, Yuki Hatanaka, Changwei Yu, Stéphane D. Vincent, Ferenc Müller, László Tora, Harry G. Leitch, Petra Hajkova, Boris Lenhard

## Abstract

Core promoters integrate regulatory inputs of genes^1–3^. Global dynamics of promoter usage can reveal systemic changes in how genomic sequence is interpreted by the cell^4^ Here we report the first analysis of promoter dynamics and code switching in the mammalian germ line, characterising the full cycle of transitions from embryonic stem cells through germline, oogenesis, and zygotic genome activation. Using Super Low Input Carrier-CAGE^5,6^ (SLIC-CAGE) we show that mouse germline development starts with the somatic promoter code, followed by a prominent switch to the maternal code during follicular oogenesis. The sequence features underlying the shift from somatic to maternal code are conserved across vertebrates, despite large differences in promoter nucleotide compositions. In addition, we show that, prior to this major shift, the promoters of gonadal germ cells diverge from the canonical somatic transcription initiation. This divergence is distinct from the promoter code used later by developing oocytes and reveals genome-wide promoter remodelling associated with alternative nucleosome positioning during early female and male germline development. Collectively, our findings establish promoter-level regulatory transitions as a central, conserved feature of the vertebrate life cycle.

## Main text

Core promoters are the sequences immediately adjacent to the transcription start site (TSS) that integrate regulatory input from enhancers and other cis regulatory elements to direct transcription initiation of genes. Selective regulation of transcription initiation defines nascent RNA production dynamics^7,8^, guides posttranscriptional control^9,10^ and can even predefine cytoplasmic RNA fate^11^. Functional diversification of general transcription factors (GTFs), which bind to the core promoter, results in distinct GTF combinations acting on subsets of genes in a cell type-specific manner (reviewed in^2,12,13^). A key outcome of this regulation is the spatio-temporal specialisation of transcription initiation of individual genes.

The nucleotide-level precision and the sequence features of transcription initiation are amongst the most informative characteristics of eukaryotic core promoters, revealing their underlying architecture and functional specialisation (reviewed in^1,3,14–16^). Analyses of transcription initiation allowed identification of distinct core promoter classes regulated by specialised factors^17^ The most prominent example is the distinction between the promoters containing TATA-boxes and, the more numerous, TATA-less promoters found in metazoan genomes. TATA-boxes (canonical TATAWAWR motif), recognised by the TATA-box binding protein (TBP), confer almost single-nucleotide precision to the initiation of transcription that occurs approximately 30 bp downstream of the start of the TATA-box (narrow promoters)^18,19^. In contrast, the majority of promoters – CpG island-overlapping TATA-less promoters – exhibit a less precise transcription initiation (broad promoters). The initiation in broad promoters occurs within a stretch of sequence serving as a “catchment area” for the transcription machinery, whose location is dependent on the stable position of the first downstream (+1) nucleosome^4,20–22^. A similar mode of transcription initiation, including core promoter sequences, has been found at enhancer regions, leading to a unified model of transcription initiation at both promoters and enhancers^23^.

Early development in zebrafish was shown to be associated with a genome-wide shift in how promoters are read: the switch from the oocyte-specific to the somatic-specific promoter usage observed during Zygotic Genome Activation (ZGA) represents one of the most dramatic instances of transcriptional regulatory rewiring known in the animal life cycle^4^ During this transition, promoters of thousands of genes (mostly ubiquitously expressed) are read differently in the context of the maternal and zygotic expression, switching from motif-dependent (maternal) to nucleosome-dependent (somatic) TSS selection upon ZGA.

Two key questions remained unsolved. The first is evolutionary: are the observed maternal to somatic TSS transitions and the promoter codes identified in zebrafish universal among vertebrates and therefore relevant to e.g. human reproduction? The second is developmental: once somatic TSSs have been established, how is it reversed during germline development to establish oocyte-specific TSS usage? The developmental stage where this somatic-to-maternal transition occurs is unknown. These questions were experimentally inaccessible until now, as Cap Analysis Gene Expression (CAGE), the gold standard method to study transcription initiation^24^, required three orders of magnitude more RNA^25^ than is realistic to obtain from a mammalian developing germ line or postnatal oocytes.

Here we report the first characterisation of transcription initiation and promoter dynamics in the early mammalian germ line, postnatal oogenesis and early preimplantation embryo. To investigate if, when, and how the somatic-maternal-somatic transitions of transcription initiation occur, we used our recently developed Super Low-Input Carrier-CAGE (SLIC-CAGE)^5,6^ to determine single-nucleotide resolution TSSs in 12 developmental stages (Figure 1a). Our data revealed that hundreds of genes shift TSS usage within their promoters from maternal (in the oocyte) to somatic (in the early embryo and zygote); and back again from somatic (in the early embryo, primordial germ cells (PGCs) and gonadal germ cells (GGCs) to maternal (follicular stage oocyte); thus documenting the existence of a germline cycle in TSS recognition. Our findings demonstrate that this promoter code is a universal feature of the vertebrate germline. In addition to this genome-wide switch, we report for the first time a change in promoter usage that occurs in the gonadal germ cells at the start of the meiotic (female) or mitotic (male) arrest, revealing the existence of an extensive promoter architecture remodelling following epigenetic reprogramming and the onset of sex differentiation in the female and male germ lines.

**Figure 1.**
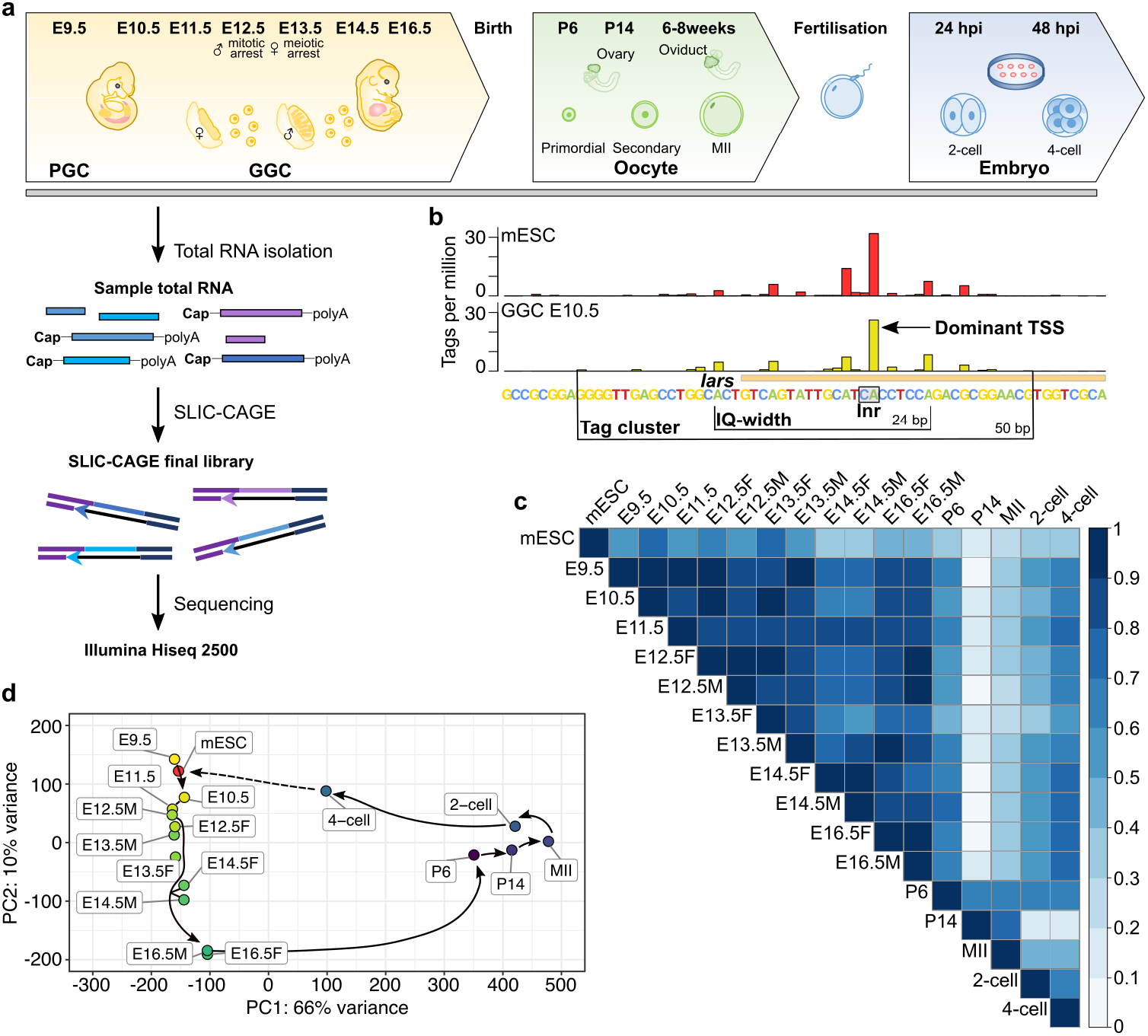
Transcription initiation at 1 bp resolution in the mouse early germ line, postnatal oogenesis and early embryogenesis. **a)** Study design – all collected stages are presented (PGC-primordial germ cell; E-embryo day; GGC – gonadal germ cell; P – postnatal day). **b)** Example CTSS signal in the isoleucyl-tRNA synthetase *(Iars)* promoter. Each bar represents one CTSS, and its height reflects the expression level. Dominant TSS, tag cluster width and IQ-width are marked. **c)** Pearson correlation of all samples at the CTSS level. **d)** Principal component analysis of stages based on consensus tag clusters expression (tag clusters aggregated from all samples into a single set). Black lines and arrows mark the differentiation time-course.

### Mapping transcriptional landscapes throughout germ line development, oogenesis and early embryogenesis reveals a promoter dynamics cycle in mice

We previously showed that SLIC-CAGE quantitatively reflects transcription and therefore can be used for joint expression profiling and promoter characterisation of scarce biological material^5^Ą We used 5-20 nanograms of total RNA from stages across mouse germline development: one stage of migratory primordial germ cells (embryonic (E) day 9.5), six stages of gonadal germ cells (E10.5 – E16.5, male (M) and female (F)), three stages of postnatal oocyte development (postnatal (P) day 6 (P6) – primordial follicle stage, P14 – secondary follicle, and ovulated meiosis II (MII) oocytes), and two early embryo stages (early 2-cell and 4-cell); and performed low-input unbiased mapping of TSSs (Figure 1a, see Methods). Regardless of the RNA input amount, we can recapitulate single-nucleotide resolution TSSs for all stages (Extended Data Figure 1a).

Tight clusters of CAGE-identified TSSs (CTSSs) originate from the same transcription preinitiation complexes, and thereby represent transcription initiation from the same core promoter (Figure 1b, Extended Data Figure 1a). The CTSS with the highest expression level within the same cluster (referred to as tag cluster) is the *dominant* CTSS^26^ (Figure 1b). We constructed tag clusters by joining neighbouring CTSSs separated by 20 or fewer base pairs (bp)^26^. The genomic locations of identified tag clusters and the distributions of their interquantile widths (IQ-width; a stable measure of promoter width, Figure 1b, see Methods) indicated high quality datasets (Extended Data Figure 2a-d;^5^). The exception are the migratory PGCs E9.5, for which the RNA quantity was at the border of SLIC-CAGE detection and which showed characteristic features of a lower complexity library, such as artificially narrow tag clusters (Extended Data Figure 2a,b). The obtained SLIC-CAGE signal across stages (Figure 1a) allows the discovery of 1) TSSs at 1 bp resolution; 2) TSS usage dynamics; 3) changes in transcription initiation precision and positional preference, alongside 4) the quantitative measurement of transcription (Figure 1b, Extended Data Figure 1a-c)^6^. Genome-wide correlation of CTSS expression confirmed a clear separation of germ cells, oocytes and early embryos (Figure 1c).

The full cycle of promoter activity along the differentiation time-course was recapitulated by principal component analysis (PCA) of both tag cluster (promoter) expression levels (Figure 1d) and CTSS (all CTSSs, Extended Data Figure 2e,f), with late GGCs (E16.5F/M) clearly segregating from the earlier stages. The early 2-cell embryo grouped closely with the oocytes, indicating that the majority of transcripts are still of maternal origin (deposited in the oocyte), while the 4-cell embryo moved halfway to mESC cells along the PC1 component, in line with the timing of the major ZGA in the mouse and the time needed to replace the maternal with the zygotic transcriptome^27,28^. Taken together, these results show that both promoter and CTSS level expressions can recapitulate developmental trajectories and support the high quality of the obtained SLIC-CAGE datasets.

### Mouse ZGA is accompanied by a shift from the maternal to somatic TSS selection rules that are conserved across vertebrates

A close examination of TSS dynamics revealed genome-wide shifts of TSSs within the same promoters (Figure 2a). To further explore this, we directly compared TSSs before and after mouse ZGA, i.e. in MII oocyte (maternal) and 4-cell embryos (somatic). We identified a total of 851 MII oocyte vs 4-cell embryo ‘shifting’ promoters – promoters exhibiting a significant change in the CTSS usage distribution or in the position of the dominant CTSS between the MII oocyte and the 4-cell embryo (Extended Data Table 1, Supplementary Table 1). These shifts indicate that the TSS selection in the oocyte is guided by rules distinct from those utilised in the early embryo, i.e. maternal vs somatic TSS usage “grammar”. We note that the TSS shifts identified using SLIC-CAGE are TSS-redistributions within the same promoter region and do not correspond to the alternative maternal promoters reported to be used by the oocyte^29^. On the contrary, SLIC-CAGE-identified oocyte tag clusters are on average closer to the UCSC-based mm10 annotation than the alternative promoter locations inferred from RNA-seq in Veselovska et al., many of which lack CAGE evidence (Extended Data Figure 3). Our CAGE-identified promoters display a consistency in sequence properties within both somatic and maternal promoters, as well as systematic differences between the two (see below), providing the most accurate single-nucleotide resolution, TSS-based annotation of oocyte transcripts.

**Figure 2.**
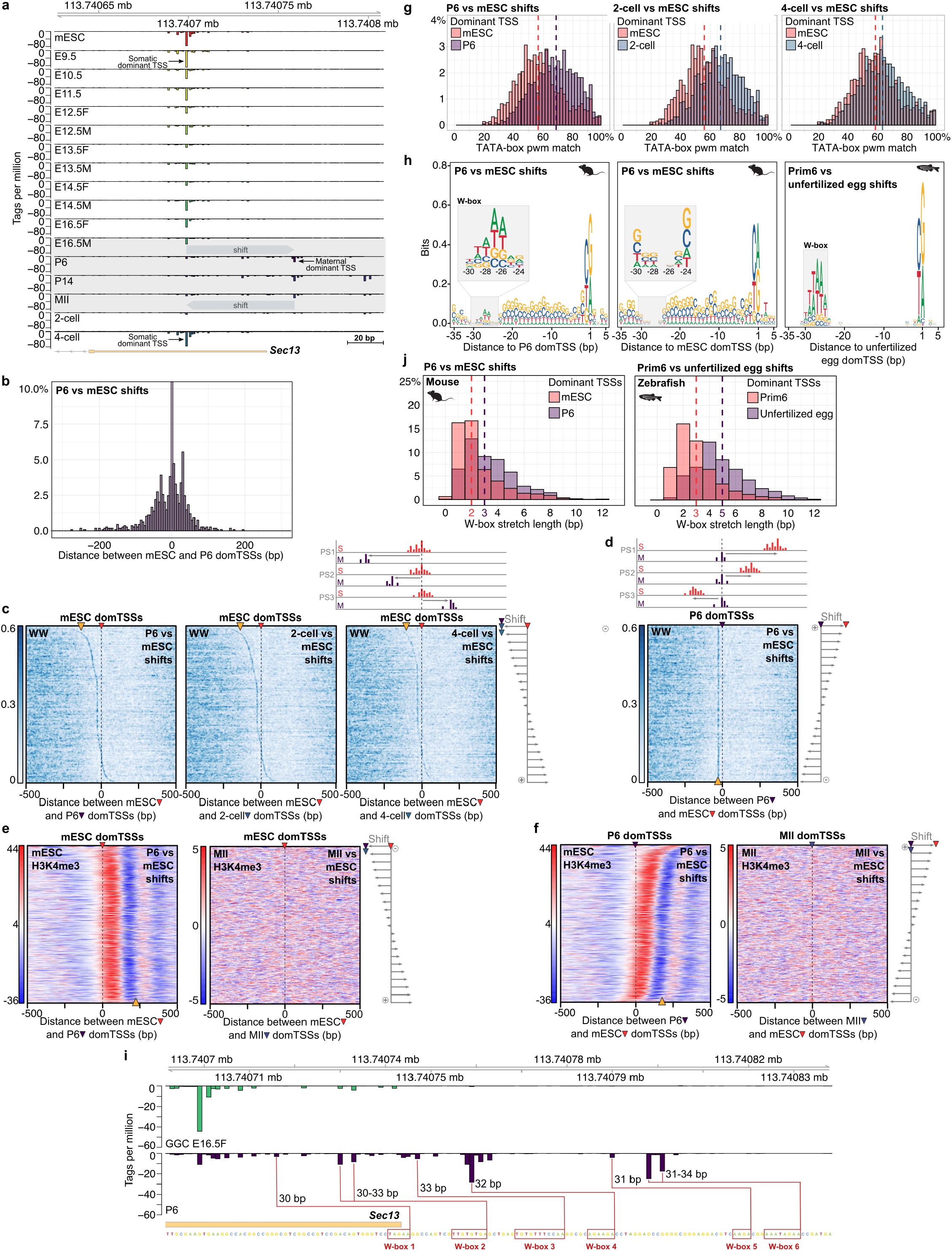
Somatic-maternal-somatic shifts of transcription initiation. **a)** CTSS signal of an example shifting promoter – *Sec13* gene. **b)** Distribution of distances between P6 and mESC dominant TSSs. **c)** WW density (top heatmaps) in shifting promoters centred to the mESC dominant TSSs (marked with a red arrow) and sorted by the distance and orientation of the shift (scheme on the top and right). Orange arrowheads indicate WW enrichment. Scale represents the WW density. **d)** P6 vs mESC shifting promoters as in c) but centred on the P6 dominant TSSs. **e)** subtracted H3K4me3 coverage (mESC or MII) of reads mapping to the plus (shown in red), and minus strand (shown in blue), centred to mESC (red arrowhead) dominant TSSs (schematics explaining H3K4me3 heatmaps is in Extended Data Figure 3c). Orange arrowhead at the bottom of the H3K4me3 heatmap marks the internucleosomal regions. **f)** Same as in e), but centred on the P6 (purple arrowhead) or MII (blue/purple arrowhead) dominant TSSs. **g)** Distributions of TATA-box pwm percentile matches in shifting promoters, centred on the sample or mESC dominant TSSs. TATA-box matches are calculated in a 1 bp sliding window encompassing 35 to 21 bp upstream of the dominant TSSs. The highest score per each sequence is reported. Vertical dashed lines mark the median. **h)** Sequence logos encompassing 35 bp upstream and 5 bp downstream of the dominant TSSs in P6 vs mESC shifts centred on the P6 dominant (left), mESC dominant TSSs (middle) or zebrafish shifting promoters (right) centred on the maternal (unfertilised oocyte) dominant TSSs. **i)** CTSS signal in the promoter region of the *Sec13* gene shown in a). Sequence is shown below the CTSS signal and the putative W-boxes driving transcription initiation in the P6 oocyte are marked in red. Distances from CTSSs and W-boxes are highlighted. **j)** Length distribution of W-box stretches identified in mouse P6 vs mESC shifting promoters (top panel), 34 to 23 bp upstream of P6 (maternal) or mESC (somatic) dominant TSSs and in zebrafish shifting promoters (bottom panel), 34 to 23 bp upstream of the maternal (unfertilized egg) or somatic dominant TSSs (prim6). The longest W-box stretch in each region is selected. Vertical dashed lines represent the median W-box stretch length.

To examine closely what happens in the oocytes and early embryos, we compared separately the TSSs from P6, P14, MII oocytes and early 2-cell, 4-cell embryos with the TSSs identified in mouse embryonic stem cells (mESC). We used mESCs as a reference because 1) the TSS signal was obtained using an equivalent, but non-low-input CAGE; 2) mESCs are sequenced at a depth comparable to our SLIC-CAGE samples; and 3) there are no significant differences in the TSS initiation between mouse somatic tissues and mESCs^20^. Therefore, herein we refer to mESC transcription initiation as an example of somatic transcription initiation. Using this approach, we identified 1098 P6-, 1130 P14- and 1028 MII oocyte shifting promoters, along with 1278 early 2-cell and 1198 4-cell embryo shifting promoters, i.e. promoters exhibiting a shift or a redistribution of CTSSs compared to mESC CTSSs, indicating a shift in the TSS grammar used (referred further as sample vs mESC shifting promoters or shifts, Extended Data Table 1, Supplementary Table 2). GO analyses of these promoters (Extended Data Figure 4a,b) revealed housekeeping terms, indicating that housekeeping genes require both TSS grammars to remain transcribed in all cell types. As housekeeping genes, these shifting promoters are GC- and CpG rich, and higher expressed than average (Extended Data Figure 4c,d). The sets of promoters identified as shifting between each oocyte stage and mESCs were highly overlapping (Extended Data Figure 4e), therefore we focused on the P6 vs mESC shifts for subsequent analyses. The physical shifts between the P6 and the mESC dominant TSSs occurred almost symmetrically in either direction, with a slight preference for the larger shifts upstream of the mESC dominant TSSs (Figure 2b), similarly to other shifting sets (Extended Data Figure 4f).

These promoter usage shifts were reminiscent of the TSS shifts between zebrafish maternal and somatic initiation^4^, suggesting a conserved mechanism across vertebrates. In zebrafish, the key identified determinants of maternal transcription initiation grammar are degenerate TATA-like motifs called W-boxes. However, mammalian housekeeping promoters are considerably more GC- and CpG-rich than their zebrafish counterparts, which could affect how often the TATA-like W-boxes appear at random. We therefore asked if the W-boxes are conserved across vertebrates, or if they diverged along with the overall promoter sequence composition. To this end, we visualised the WW (A,T/A,T) dinucleotide density in the shifting sequences aligned to the mESC dominant TSSs and sorted by the distance and the orientation of the shift between the corresponding dominant TSSs (Figure 2c, top right scheme, Extended Data Figure 5a,b). A strong WW enrichment aligned with the positions of the P6 oocyte and early 2-cell embryo dominant TSSs, implying that its selection is guided by the TATA-like elements (Figure 2d, see schemes in Extended Data Figure 6a,c). The WW enrichment decreased in the 4-cell embryo, indicating that the underlying shift is a leftover from the maternal/oocyte rules of transcription initiation, in line with the timing of the mouse major ZGA. We further quantified this by matching the regions 35 to 21 bp upstream of the dominant TSSs to a TATA-box position weight matrix (pwm), used as a proxy for TATA-like elements (Figure 2g, see Methods). All shifting promoters when centred to the P6, 2-cell or 4-cell embryo dominant TSSs exhibited higher TATA-box pwm scores than when centred to the mESC dominant TSSs, demonstrating a higher usage of the TATA-like elements in the maternal grammar. This is weaker in the 4-cell than the 2-cell embryo, revealing again a transition from the maternal transcription initiation code.

It was shown previously that the position of the dominant TSSs in broad promoters occurs at a fixed distance from the first downstream (+1) nucleosome, about 125 bp upstream of the nucleosome dyad^4,21,22^. In agreement with this, when expressed in mESCs, the TSSs used by the P6 vs mESC shifting promoters showed a clear alignment with the H3K4me3-marked +1 nucleosomes (mESC H3K4me3 ChIP-seq), indicating that its selection depends on the position of the +1 nucleosome (Figure 2e, Extended Data Figure 5b,d,e, for schemes see Extended Data Figure 5c and 6b,d). In contrast, there was no clear H3K4me3-marked +1 nucleosome alignment to the MII oocyte dominant TSSs when compared with MII H3K4me3 ChIP-seq data (Figure 2e), as expected for promoters with motif-dependent TSS selection^4,21,22^.

Sequence composition of the shifting promoters revealed an enrichment of the TATA-like elements approximately 30 bp upstream of the primordial oocyte (P6) dominant TSSs (Figure 2h, left panel). In contrast, these TATA-like elements do not occur at a fixed position upstream of the mESC dominant TSSs (Figure 2h, middle panel). The motif and tetranucleotide composition analysis of these TATA-like elements revealed that in a large majority of cases, they do not correspond to a canonical TATA-box, but rather to a more degenerate TATA-like W-box motif (Figure 2h, left panel, inset, Extended Data Figure 7). Henceforth we refer to this TATA-like element as a W-box. We also note that there can be more than one W-box element per promoter, each with a prominent CTSS about 30 bp downstream of it, leading to a composite broad transcription initiation pattern (Figure 2i). This is distinct from the somatic, +1 nucleosome dependent pattern, establishing the multiple W-box architecture as a new promoter class. The relative locations of these W-boxes with respect to the nucleosome positioning signal determine if there is a shift of the maternal dominant TSSs, and whether this is upstream or downstream of the somatic +1 nucleosome-determined dominant TSS. This maternal promoter architecture (single and multiple W-boxes) suggests that the overlap of the somatic and maternal promoter codes and their sequence properties are a universal property of vertebrate genomes^4^. Intriguingly, these features have been preserved despite large differences in the GC content of the sequences in which they are embedded in each genome (zebrafish – 36.9%, mouse – 49.8 % in promoters, see Methods. Figure 2h, left vs right panel). A measurable effect of the increased GC content on the mouse W-boxes is that the stretch of As and Ts is on average shorter in the mouse (median 3 in P6 oocyte vs mESC shifts; Figure 2j, left) than in zebrafish (median 5 in Prim6 post-ZGA embryo vs unfertilized egg shifts; Figure 2j, right). While the zebrafish W-boxes typically consist of 4-5 As and Ts, in mouse, a stretch of two or three A/Ts embedded in the GC-rich sequence can serve the same function (Figure 2h).

To examine if the change in grammar occurs universally in all promoters, not only in those in which the CTSSs shift, we performed an unbiased analysis of all promoters active in all samples, both maternal and zygotic, not requiring a displacement of CTSSs. We hypothesized that the same promoters can have an overlapping grammar (maternal W-box and somatic grammar), reflected by different TSS distributions, even if the dominant TSS does not change. Using promoter-annotated tag clusters, we identified a set of 6964 ubiquitously expressed promoters (cluster 1, Extended Data Figure 8a,b,d). These ubiquitously expressed genes used more W-box elements in the oocytes and the early 2-cell embryo than in mESCs (Figure 3a, Extended Data Figure 9a). Further examination of this set showed a stronger enrichment of the W-box elements 30 bp upstream of the maternal dominant TSSs, no precise +1 nucleosome positioning downstream of the maternal dominant TSSs (H3K4me3-marked, MII oocyte data), and precisely ordered +1 nucleosomes (H3K4me3-marked, mESC data) downstream of the somatic dominant TSSs in mESCs (Figure 3b). H3K4me3-marked +1 nucleosomes in mESCs (H3K4me3-marked mESC) align with a large subset of maternal dominant TSSs, as the majority of ubiquitously expressed promoters does not change the dominant TSS positions when transitioning from the somatic to maternal rules of transcription initiation (ubiquitously expressed genes (SOM 1, 6964 promoters), overlap with 1035 P6-, 1094 P14 and 983 MII-shifting promoters, i.e. about 15% of ubiquitously expressed promoters change the dominant TSS position). In line with the housekeeping function, expression of the ubiquitous promoters is largely unchanged by the shift from the somatic to maternal grammar (Extended Data Figure 8c, Extended Data Figure 9b).

**Figure 3.**
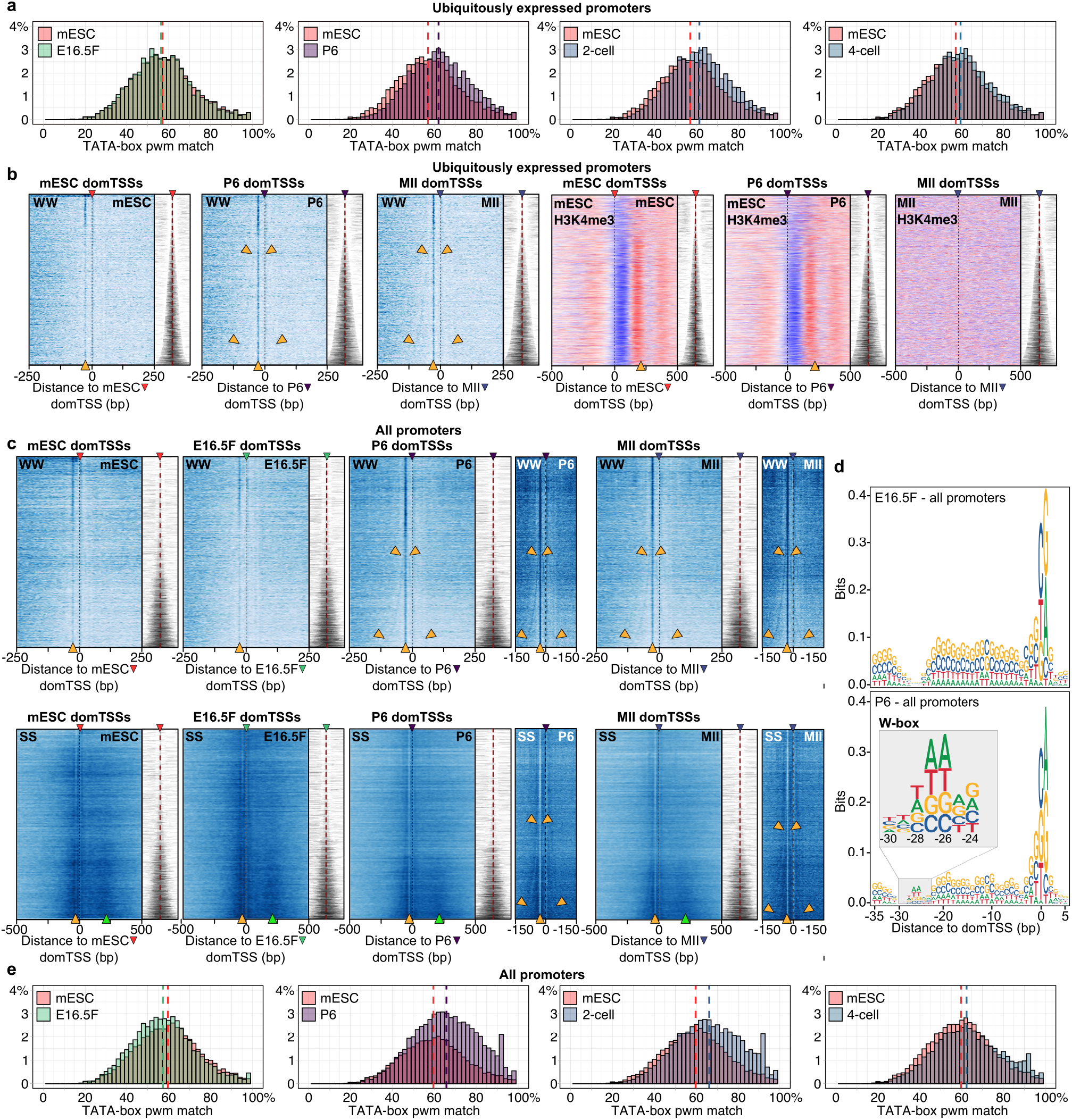
Ubiquitously expressed genes use both somatic and maternal grammar, established in the primordial (P6) oocyte. **a)** Distributions of TATA-box pwm percentile matches in ubiquitously expressed tag clusters (SOM 1, see Extended Data Figure 8). TATA-box pwm percentile matches are calculated in a 1 bp sliding window encompassing a region 35 to 21 bp upstream of the dominant TSSs. Only best match per sequence is reported. Vertical dashed lines mark the median. **b)** Heatmaps showing WW density and mESC H3K4me3 in ubiquitously expressed genes (SOM cluster 1), centred on the sample specific dominant TSSs (labelled on top of heatmaps with an arrowhead in a sample-specific colour – see colour scheme used in Extended Data Figure 1a) and ordered by the IQ-width (narrower on top, broader on the bottom), shown as tag cluster coverage signal in grey on the right of each heatmap. The vertical dashed lines – black or red, in all heatmaps mark the dominant TSS positions. Orange arrowheads in WW heatmaps (bottom) mark the expected position of the WW enrichment – 30 bp upstream of the dominant TSSs, and within the P6 and MII heatmaps also mark the outmost W-box of the multiple W-box architectures. Orange arrowheads in the H3K4me3 heatmaps mark the internucleosomal region. **c)** Heatmaps showing WW (top) or SS density (bottom) in all mESC, E16.5F GGC, P6 oocyte or 4-cell embryo identified promoters. Sequences are centred on the sample specific dominant TSSs (marked with an arrowhead on top of the heatmap) and sorted by the tag cluster IQ-width (shown in grey on the right, as in b). Tag cluster coverage reflecting IQ-widths is shown on the right of each heatmap. Orange arrowheads mark the WW enriched TATA-like region (WW heatmaps) or the CpG/GC rich internucleosomal region (H3K4me3 heatmaps). The orange arrowheads within the P6 and MII oocyte WW heatmaps mark the outmost W-box of the multiple W-box architectures. The narrow WW or SS heatmaps next to the P6 and MII WW density heatmaps show the region with the outer W-boxes on a different scale, to enhance its visibility. **d)** Sequence logos encompassing 35 bp upstream and 5 bp downstream of the dominant TSSs of all tag clusters in the E16.5F (top) or P6 oocyte (bottom). **e)** Distributions of TATA-box pwm percentile matches in all SLIC-CAGE identified tag clusters. Calculation is the same as in a). Vertical dashed lines represent the median.

The major ZGA occurs in the mouse at the 2-cell stage^27,28,30^. To investigate the differences of transcription initiation rules and to separate the new zygotic transcripts from the maternally inherited RNAs, we compared promoters of genes expressed exclusively or at much higher level in the embryo relative to the oocyte, and those expressed exclusively or predominantly in the oocyte. We identified these transcripts, resulting with 2631 oocyte- and 1309 embryo-specific promoters (cluster 4 and cluster 8, Extended Data Figure 8a). The embryo-specific genes have the highest expression in the 4-cell embryo stage, as the early 2-cell embryo is still largely composed of inherited maternal transcripts. GO analysis of the embryo-specific genes confirmed involvement in the early embryonic processes, while the oocyte-specific genes showed oocyte-related terms (Extended Data Figure 8e,g), and late gonadal germ cell-specific genes (cluster 5, Extended Data Figure 8a) included meiotic markers (driven by the female gonadal germ cells entering meiotic arrest (Extended Data Figure 8f)). Sequence analysis of the embryo-specific promoters revealed that the W-box elements are not enriched upstream of their dominant TSSs, unlike the oocyte-specific promoters (matches in the oocyte-specific genes are shifted towards higher percentile values, resulting in a higher median percentile match, Extended Data Figure 9c-e). Instead, these promoters are dependent on the precise +1 nucleosome positioning (Extended Data Figure 9f,g). To the contrary, the dependence of transcriptional initiation on W-box elements holds true for all promoters in the postnatal oocyte, not only the housekeeping ones with two overlapping grammars, thereby establishing the W-box as a key core promoter element in the oocytes (Figure 3b,c, Extended Data Figure 9d, Extended Data Figure 10).

### Maternal TSS grammar is established during germline development in the primordial follicle (P6) oocytes

While the shift from the maternal to somatic TSS usage occurs following the ZGA, it is unknown when the maternal TSS usage is established in the context of germline development. We thus analysed the TSS usage across our datasets (Figure 1a) for their association with W-box elements: the rationale was that the bulk transition to motif-dependent transcription initiation should indicate the stage in which the maternal grammar takes over from somatic.

We constructed heatmaps to visualise WW or SS (C,G/C,G) dinucleotide density in 500 or 1000 bp sequence windows centred on the dominant TSSs (Figure 3c) and ordered by the IQ-width of the corresponding tag cluster. PGCs (E9.5) and all gonadal germ cells (E10.5-E16.5F/M) exhibited patterns similar to mESCs, indicating their reliance on the canonical somatic promoter architecture (Figure 3c, Extended Data Figure 10a). In this architecture, only the narrow promoters (top of the heatmaps) were associated with a WW enrichment 30 bp upstream of the dominant TSSs, characteristic of the TATA-box elements (Figure 3c, Extended Data Figure 10a, orange arrowhead at the bottom of heatmaps). This WW enrichment within the narrow promoters showed a strong presence of the consensus TATAWAWR TATA-box motif, expected for the somatic type of precise transcription initiation (Extended Data Figure 10b). The broader primordial (PGC) and gonadal germ cell (GGC) promoters showed a position-specific SS enrichment (Figure 3c, marked with a green arrowhead at the bottom), typically associated with the precisely positioned +1 nucleosome^4,14,21^.

In contrast, all oocyte stages exhibited a strong WW enrichment ~30bp upstream of the dominant TSSs, independent of the promoter width (Figure 3c, Extended Data Figure 10a). Association of the WW enrichment with the “broad” maternal promoters is in accordance with the multiple W-box architecture (Figure 2i), further supported by the weak WW enrichment representing the outermost W-boxes (Figure 3b,c, see orange arrows within heatmaps). Sequence analysis of the WW enriched regions revealed a presence of the W-box elements at a fixed distance from the dominant CTSSs in the P6 oocyte, and their absence in the gonadal germ cells (Figure 3d). To quantify the usage of W-box elements, we extracted sequences from positions 35 to 21 bp upstream of the dominant TSSs of all promoters, scored them with a TATA-box pwm in a sliding window, selected the highest percentile match per sequence and visualised the distributions of the matches (Figure 3e). E16.5 GGCs (M and F) showed similar matches as mESCs, while the distribution of the best percentile matches in the P6 oocyte was shifted towards a higher percentile, in line with the WW enrichment and transition to the maternal code. Therefore, there is no evidence of maternal code usage in the GGCs, while the maternal code is fully active by the oocyte primordial follicle P6 stage.

Finally, to better determine the timing of transition to the maternal code and examine if it closely precedes the P6 stage, we searched for evidence of the residual somatic TSS signal in the P6 oocyte. We classified dominant CTSSs and selected somatic-specific dominant TSSs, i.e. dominant TSSs expressed exclusively or at a much higher level in mESC, PGC and GGCs relative to the oocyte (cluster 17 and 18, Extended Data Figure 15a). These somatic dominant TSSs are systematically higher expressed in the P6 than in the P14 oocyte (Extended Data Figure 11a, b), suggesting that the transition to the maternal type of TSS usage has just finished. We also directly compared transcription initiation in the primordial P6 and the secondary follicle P14 oocyte and identified only 23 shifting promoters. Visual inspection of these shifting promoters revealed some transcription initiation patterns in the P6 oocyte, reminiscent of the patterns observed in the early 2-cell embryo following the major ZGA, and indicating a transitional pattern (Extended Data Figure 11c-g). Overall, this low number of promoter shifts identified between the P6 and P14 oocyte, and systematically higher expression of the somatic dominant TSSs in the P6 relative to the P14 oocyte strongly suggest that the transition to the maternal type of TSS usage has recently finished, i.e. the transition occurs at, or just before the primordial follicle (P6) stage of oocyte development.

### A novel genome-wide remodelling of the somatic TSS initiation occurs in the late gonadal germ cells and precedes the transition to the maternal code

Mouse PGCs are specified in the developing embryo around embryonic day 6.25. Following the migration into developing gonads, germ cells undergo extensive epigenetic reprogramming around E11.5-E12.5, preceding sex specification and meiotic entry of the female GGCs^31,32^. As shown above, none of these events are accompanied by a significant departure from the somatic promoter code: the transcription initiation rules remain the same as in mESC.

Surprisingly however, the promoters in both male and female E16.5 GGCs showed shifts in CTSS patterns compared to the earlier GCCs, PGCs and mESCs that do not correspond to the switch to maternal code (Figure 4a,b, 829 E16.5F vs mESC and 707 E16.5M vs mESC shifting promoters; Extended Data Table 1, Supplementary Table 2). GO enrichment analyses of these shifting promoters revealed housekeeping terms (Extended Data Figure 12a,b). Promoters undergoing these CTSS shifts in gonadal germ cells are also GC + CpG rich and higher expressed than the average promoter (Extended Data Figure 12c,d). For some promoters the evidence of the beginning of this shift appears even at earlier time points (181 shifting promoters common to E14.5 vs mESC and E16.5 vs mESC GGCs, Extended Data Figure 12e). The physical shifts in the dominant TSSs occurred in both directions, with a slight preference upstream of the dominant TSSs observed in mESC (Figure 4c, 8 bp median upstream shift in dominant TSS position).

**Figure 4.**
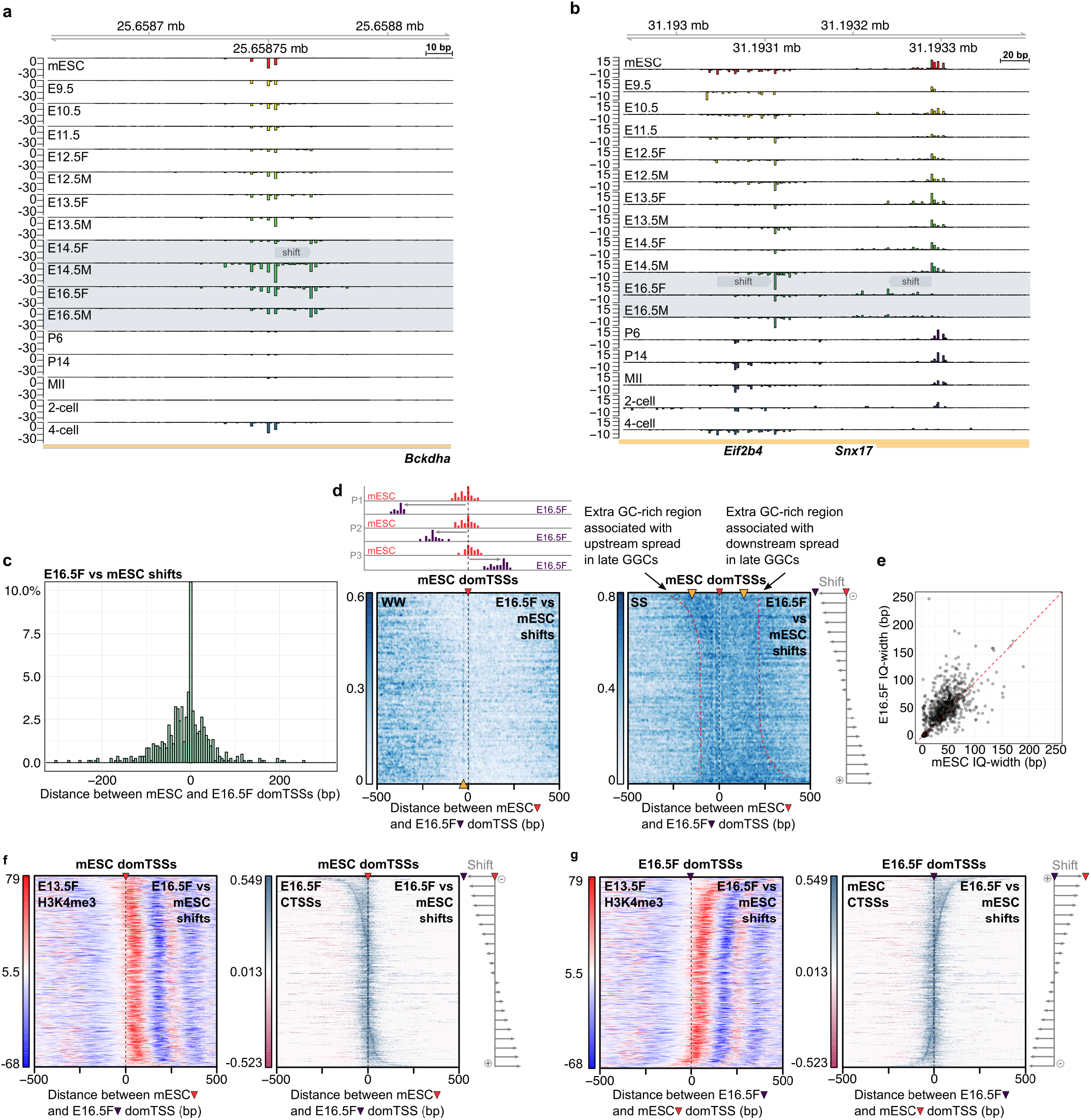
A novel genome-wide change of the somatic promoter code in the late gonadal germ cells precedes the transition to the maternal code. **a)** CTSS signal of an example shift – *Bckdha g*ene. **b)** CTSS signal of an example shifting bidirectional promoter, with broadening *(Snx17* gene) or narrowing *(Eif2b4* gene) of the signal. **c)** Distribution of the distances between the E16.5F and mESC dominant TSSs in E16.5F vs mESC shifting promoters. **d)** WW (left) and SS (right) density in E16.5F vs mESC shifts, centred to the mESC dominant TSSs (marked with a red arrowhead on top) and ordered by the distance and orientation of the shift (schemes on the top and right). Red dashed lines follow the GC enrichment. Orange arrowhead in WW heatmaps indicates the expected position of the WW enrichment (TATA-box or W-box), while in SS heatmaps they indicate regions of SS enrichment. **e)** Correlation of sample specific IQ-widths in E16.5F vs mESC shifting promoters. **f)** Heatmaps showing E13.5F H3K4me3 (left) or E16.5F CTSS coverage signal (right) in E16.5F vs mESC shifts centred on the mESC dominant TSS (marked with a red arrowhead on top) and ordered by the distance and orientation of the shift (scheme on the right of CTSS heatmap). **g)** Same as in f) albeit centred on the E16.5F dominant CTSSs and showing mESC CTSS coverage signal.

The E16.5F vs mESC shifting promoters show an overall enrichment in the GC content in the direction of the shift (Figure 4d, right). These promoters also: 1) lack the WW enrichment 30 bp upstream of the dominant CTSSs characteristic of the oocyte vs mESC shifting promoters (Figure 4d, left); 2) are mostly broad (median IQ-width is 49 bp), and 3) the majority (62 %) were broader in the E16.5F sample than mESCs (Figure 4e), without a significant change in the expression level (Extended Data Figure 12g).

The dominant TSS position of a broad promoter in somatic cells mainly depends on the precise +1 nucleosome localisation^14,20,21^. The broadening of the TSS signal in the E16.5 GGCs suggests that either there are changes in the underlying +1 nucleosome position, or an unknown mechanism extends the “catchment area” within which transcription can initiate from the unchanged +1 nucleosome position, thereby resulting in a broader promoter architecture. To distinguish between those two scenarios, we compared the strength of the WW periodicity signal downstream of the dominant TSS – a clear indication of precise +1 nucleosome positioning, in the E16.5F vs mESC shifting promoters (Extended Data Figure 12j). The WW periodicity signal in the E16.5F vs mESC shifts was somewhat stronger when centred to the E16.5F dominant TSSs compared to a TSSs randomly selected within the same tag cluster (Extended Data Figure 12k, see further support below and in Extended Data 12l,m).

To test if the underlying sequence of the GGC shifting promoters is associated with a switch to an alternative nucleosome positioning signal, we produced a pwm representation of the nucleosome-preceding sequence motif (Extended Data Figure 13a,b, see Methods). In all cases, regardless of the centring point (E16.5F or mESC dominant TSSs), there was a match peak aligned with the dominant TSS (Extended Data Figure 14c,d), suggesting that the extra GC-rich sequence seen in Figure 4d accommodates this signal. This supports the model in which the changes in the E16.5 TSS patterns are associated with an alternative sequence-encoded nucleosome positioning signal becoming accessible to the transcription initiation machinery. A robust follow-up analysis of the autocorrelation of the WW density signal (Extended Data Figure 12l,m), high correlation of the physical shifts in tag cluster borders (Pearson correlation = 0.93, Extended Data Figure 13e), and a detailed analysis of the GGC specific dominant TSSs (Extended Data Figure 15) further support the alternative nucleosome positioning model.

We sought to verify our sequence-based model predictions using experimental data reflecting nucleosome positions. We used the H3K4me3 ChIP-seq data from E13.5F and E17.5M GGC stages because 1) only these were of sufficient quality to allow the visualisation of +1 nucleosome; 2) SLIC-CAGE monitors accumulated RNA, so changes in the TSS selection may reflect a change in the nucleosome configuration from an earlier stage; 3) The onset of the promoter shifts in GGCs can be observed in earlier stages (Figure 4a, Extended Data Figure 12e), and the classification of the dominant CTSSs showed that some GGC-specific dominant TSSs start being expressed in the 13.5F stage (Extended Data Figure 15a, clusters 22 and 24); and 4) GGC shifts occur in both female and male germline. We centred the E16.5F vs mESC shifting promoters on the mESC or E16.5F dominant CTSSs, ordered them by the distance and orientation of the shift, and visualised H3K4me3 signal from E13.5F and E17.5M stages (Figure 4f,g, Extended Data Figure 12n). Remarkably, both mESC and E16.5F CTSS signals correlate well with the nucleosome-free region, while the +1 nucleosome signal is not as precisely aligned with the mESC dominant TSS, as expected for the somatic nucleosome-dependent transcription initiation (Figure 4f,g). This provides strong evidence to support our hypothesis that nucleosomes do shift. Similar was observed using the E17.5M H3K4me3 data (Extended Data Figure 12n). Taken together, our analyses revealed a novel somatic transcription initiation rule in the late GGCs, where the alternative +1 nucleosome positions guide TSS selection.

Finally, to examine what happens with E16.5F vs mESC shifting promoters in the oocyte context, we divided them into those that shift only compared to mESC (single) or shift twice – compared to the mESC and compared to the oocyte (double shift, Extended Data Figure 14). We show that both single or double shift are equally likely, and that transcription initiation in the late GGCs in both classes shows dependence on the +1 nucleosome positioning, regardless of the fate in the oocyte. This indicates that the late GGC and oocyte shifts are architecturally and functionally independent.

## Discussion

Our study reveals systemic changes in the rules of transcription initiation in the mouse life cycle, manifested through physical shifts in the TSS usage (Figure 5). Maternal/oocyte-specific transcription relies on the motif-dependent transcription initiation at fixed distances downstream of the weak TATA-like elements termed W-boxes, similarly as shown for zebrafish^4^ Multiple W-boxes in a single promoter lead to a broad transcription initiation pattern which is a superposition of multiple narrow initiation points, each driven by its own W-box about 30 bp upstream. This architecture is distinct from the canonical somatic broad type, establishing this maternal code as a novel motif-dependent broad promoter class. We further show that during germline development, the shift to maternal W-box dependent transcription initiation occurs just before the primordial follicle (P6) oocyte stage. W-box sequences are functionally equivalent to the TATA-box sequence, suggesting that transcription initiation in the oocyte may be mediated by TBPL2, shown to replace the general transcription factor TATA binding protein (TBP) in the growing oocyte^33,34^ (Extended Data Figure 16a). Indeed, we recently demonstrated^35^ that the non-canonical TBPL2/TFIIA complex replaces the TFIID (TBP/TAFs) in the growing oocyte, and that the deletion of *Tbpl2* causes genomewide changes in the TSS selection of the secondary follicle (P14) oocyte, evidenced also by a decrease in the usage of the W-box elements for TSS selection. These results strongly support the notion that the shifts in the TSS selection identified already in the primordial follicle (P6) oocyte are caused by a global transition to the maternal W-box dependent grammar, read by an oocyte-specific transcription machinery TBPL2/TFIIA. Since the TBPL2 orthologs separate from TBP are found only in jawed vertebrates, this suggests that its specialisation has arisen through one of two basal rounds of vertebrate whole-genome duplication.

**Figure 5.**
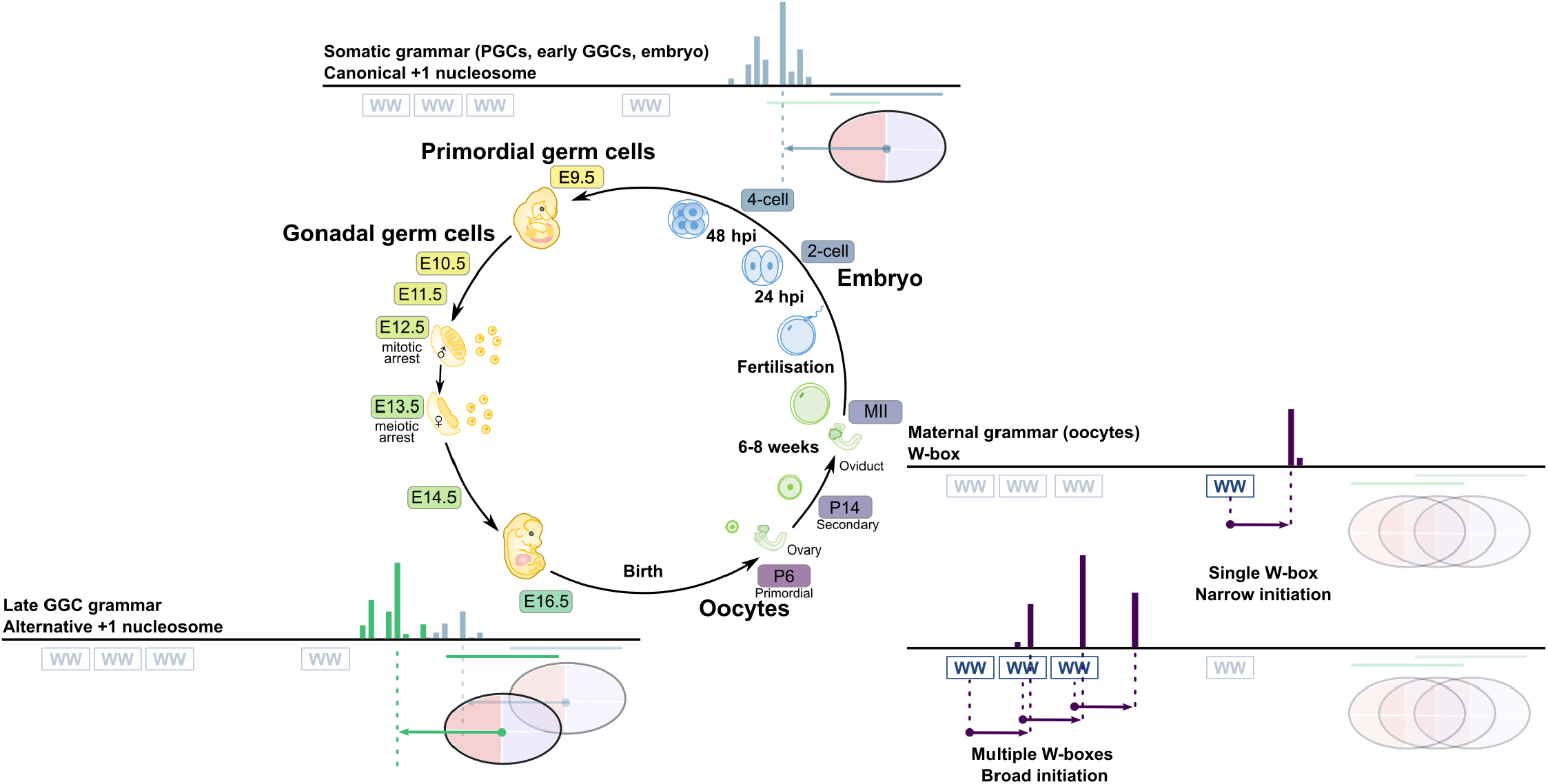
Canonical somatic +1 nucleosome-dependent, alternative +1 nucleosomedependent and maternal transcription initiation grammar. Somatic canonical +1 nucleosome-dependent grammar is used in primordial germ cells, early gonadal germ cells and embryo upon zygotic genome activation. Alternative +1 nucleosome-dependent grammar is used in late gonadal germ cells. Maternal W-box grammar is used in oocytes. Two promoter architecture classes are shown – single W-box leading to a narrow transcription initiation, and multiple W-boxes leading to a broad transcription initiation pattern, distinct from the +1 nucleosome somatic type.

Both PGCs and GGCs exhibit canonical somatic rules of transcription initiation, as no dramatic changes in the core promoter usage are detectable compared to the mESC (used in this study as a reference for canonical somatic transcription initiation). This demonstrates that globally, the somatic type of transcriptional initiation can sustain genome-wide epigenetic reprogramming occurring in the migratory PGCs and later during the early gonadal stages (E10.5-E11.5). Intriguingly, in the late gonadal germ cells, hundreds of promoters exhibit a shift in the TSS selection unrelated to that of the subsequent maternal promoter code. These shifts predominantly lead to broadening of the promoters, suggestive of changes in the nucleosome configuration and the associated TSS “catchment area” of promoters with nucleosome-dependent TSS selection (Figure 5).

What exactly increases the accessibility of these regions remains unknown, but SLIC-CAGE provides a nucleosome-resolution monitoring method for future mechanistic studies. Additional features that may influence nucleosome reconfigurations are histone composition of the +1 nucleosomes and alternative general transcription factors. Non-canonical histone variant H3.3 is encoded by *H3f3a* and *H3f3b* in the mouse genome and it is incorporated into chromatin in transcriptionally active areas that are undergoing nucleosome displacement (reviewed in (Szenker et al. 2011)). As expected for non-proliferative cells, we see a significant increase in the expression levels of both gene variants in the male and female late gonadal germ cells (Extended Data Figure 16b). Interestingly, the E16.5F vs mESC shifting genes show a significant decrease in expression levels upon H3.3 knockdown (Extended Data Figure 16c), supporting the hypothesis that H3.3 may be involved in facilitating nucleosome reorganisation and expression of this gene set. In terms of alternative general transcription factors, a recent study showed that TFIID alternative subunits *Taf4b, Taf7l* and *Taf9b* are co-ordinately upregulated during germline development (Gura et al. 2020). This trend is recapitulated in our SLIC-CAGE time-course data (Extended Data Figure 16d). Further investigation will be needed to distinguish if changes in nucleosome composition, general transcription factor composition, or both, facilitate nucleosome reconfigurations in late GGCs.

A fundamental biological question is – what problem does the change in transcription initiation rules in the growing oocyte solve? The most obvious hypothesis is that this mechanism can efficiently change the entire transcriptional repertoire of the cell by shutting one (somatic) mechanism at all promoters recognised by it, and replacing it with another (maternal) one. This simultaneously shuts down all promoters needed in the somatic cells only, turns on those needed only in the oocyte, and shifts the TSS pattern in thousands of genes that need to be expressed in both. Gametes are probably the only cell type in which such abrupt switch is the right solution to the problem: in all other contexts when gene regulation changes, either a limited set of genes needs to be turned on or off, or the changes need to be introduced gradually, in an orderly regulatory cascade. In addition, gametogenesis involves changes in DNA methylation, ploidy and chromatin conformation that could disrupt mechanisms ensuring gene dosage and long-range regulation, and a switch to promoters that respond to those mechanisms differently is more robust than other available mechanisms of transcriptional regulation.

Overall, our findings demonstrate that promoter transitions are a conserved feature of the vertebrate germ line, and that the genome wide changes in promoters are the mechanistic basis for the most comprehensive rewiring of expression in the vertebrate life cycle.

## Methods

### Germ cell collection

All animal experiments were carried out under and in accordance with a UK Home Office Project Licence in a Home Office-designated facility. Primordial and gonadal germ cells were isolated from embryos obtained from a 129Sv female and GOF18ΔPE-EGFP^36^ male cross. Genital ridges from 1-2 litters (7-8 embryos per litter) were dissected out and digested at 37°C for 3 min using TrypLE Express (Thermo Fisher Scientific). For E9.5 and E10.5 stages, posterior half of the embryo was dissected out and digested with trypsin solution (0.25% trypsin, 10% chick serum (both from Life Technologies), 1.27mM EDTA (Sigma)). Litters were pooled for stages up to and including E11.5 to obtain more material. Enzymatic digestion was neutralized with DMEM/F-12 (Gibco) supplemented with 15% fetal bovine serum (Gibco) after manual dissociation by gentle pipetting. The cells were spun down by centrifugation and resuspended in 0.1% BSA PBS. GFP-positive cells were isolated using an Aria Fusion (BD Bioscience) flow cytometer and sorted into ice-cold PBS. Total RNA was isolated from using DNA/RNA Duet Kit miniprep kit (Zymo Research, USA).

### Collection of Oocytes, *in vitro* Fertilization (IVF) and Embryo Culture to obtain 2-cell and 4-cell embryos

In this study, all preimplantation embryos were prepared by IVF. Collection of P6 and MII oocytes, and fertilized embryos was performed based on previous reports^37–39^. P6 oocytes were collected from C57BL/6J strain female mice at P6. The ovaries were dissected well with a microscissors before being incubated in 0.05% collagenase (gibco) dissolved in Dulbecco’s modified Eagle’s medium-F12 (DMEM/F12; Invitrogen) supplemented with 4 mg/ml bovine serum albumin (BSA; SIGMA), 80 μg/ml Kanamycin (SIGMA), with frequent pipetting. After 30 min, the oocytes were picked up using mouth pipette and transferred to PBS solution for washing. To select the oocytes at primordial follicle stage, the size of the oocytes was measured by an imaging software system (Octax EyeWare, Octax) and the 20-30 μm of oocytes were collected for further experiments. MII oocytes were collected from C57BL/6J female mice that were superovulated with serum gonadotropin (PMSG-Intervet; MSD) and human chorionic gonadotropin (hCG; CHORULON) at 46-to 52-h intervals. Spermatozoa were collected from the cauda epididymides of C57BL/6J male mice. The sperm suspension was preincubated in human tubal fluid (HTF) medium for 1.5 h at 37 °C under 5% CO2 in humidified air. The sperm suspension was added to the oocyte cultures, and morphologically normally fertilized oocytes were collected 2 h after insemination. Fertilized embryos were cultured in potassium simplex optimized medium (KSOM) at 37 °C under 5% CO2 in humidified air. Total RNA was isolated from the samples using the DNA/RNA Duet Kit miniprep kit (Zymo Research, USA).

### SLIC-CAGE library preparation

SLIC-CAGE libraries were created using the published SLIC-CAGE protocol^5,6^. The amount of RNA used in SLIC-CAGE ranged between 5-20 ng of total RNA per stage. Minimum of two biological replicates were prepared per stage. Eight samples with different barcodes were standardly multiplexed prior to sequencing. The libraries were sequenced on a HiSeq2500 in a single-end 50 bp mode (Genomics Facility, MRC, LMS). Barcodes were prior to usage confirmed not to introduce any bias by using performing technical replicates of nAnT-iCAGE protocol on *Saccharomyces cerevisiae* total RNA. All samples regardless of the barcode used showed a high correlation at the level of individual CTSSs (Pearson correlation coefficient = 0.98 – 0.99).

### Processing of SLIC-CAGE tags

Sequenced libraries were demultiplexed using CASAVA allowing zero mismatches for barcode identification. Demultiplexed CAGE tags (47 bp) were mapped to a reference mm10 genome using Bowtie2^40^ with default parameters that allow zero mismatches per seed sequence (default 22 nucleotides). The mapped reads were first sorted using samtools^41^ and only uniquely mapped reads were kept for downstream analysis in R graphical and statistical computing environment (http://www.R-project.org/). The mapped and sorted unique reads were imported into R as bam files using the standard workflow within the CAGEr package v 1.20^26^. The 5’G that is commonly added through template free activity of reverse transcriptase was removed using CAGEr G-correction workflow: 1) if the first nucleotide in a read is a G and flagged as a mismatch, it is removed from the read; 2) if the first nucleotide is G and not a mismatch, it is removed or retained according to the percentage of identified mismatched G’s. All unique 5’ends of reads are CAGE-supported transcription start sites (CTSSs) and the number of each CTSS (number of tags) reflects expression levels. The highest expressed CTSS is termed the ‘dominant CTSS’. Raw tags were normalised using a referent power-law distribution and expressed as normalized tags per million (tpm)^42^. Biological replicates were highly correlated and were therefore merged prior to downstream analyses using standard Bioconductor packages (http://www.bioconductor.org/) and custom scripts.

mESC CAGE data used in the manuscript is from E-MTAB-6519 (ArrayExpress,^5^). P14 oocyte data is from E-MTAB-8866 (ArrayExpress^35^).

### Clustering CTSSs into tag clusters

Tag clusters represent a single functional transcriptional unit, i.e. all transcription start sites where RNA polymerase II initiates and produces the same transcript. CTSSs that passed the threshold of 1 tpm in at least 1 sample were clustered together using distance-based clustering within the CAGEr package, with 20 bp allowed as maximum distances between neighbouring

CTSSs. Width of each tag cluster was calculated as interquantile width (IQ-width) to exclude effects of extreme outliers. For each tag cluster, a cumulative distribution of signal was calculated, and its boundaries expressed using the 10th and 90^th^ percentile of the signal, as implemented in CAGEr. The distance between the boundaries is the IQ-width of a tag cluster. To allow between-sample comparisons, tag clusters with sufficient support (at least 3 tpm) within 100 bp distance were aggregated across samples to define consensus clusters.

### Genomic locations of tag clusters

Genomic locations of tag clusters were determined using the ChIPseeker package^43^. Promoters were defined as 600 bp windows encompassing 500 bp upstream and 100 bp downstream of the annotated transcriptions start sites (annotations as in TxDb.Mmusculus.UCSC.mm10.knownGene).

### Sample correlations and PCA analyses

Sample correlation analyses was done using CAGEr normalised expression values of CTSSs (power-law normalisation, see above). Pearson correlations were then calculated between samples and visualized using the corrplot package (CRAN).

PCA analyses were produced using DESeq2 Bioconductor package. Either raw expression values (tpm) from CAGEr created consensus clusters were used (tag clusters aggregated from individual samples into a single set; see above Clustering CTSSs into tag clusters) or raw expression values of individual CTSSs. Expression values were normalised using the vst (variance stabilising transformation) function of the DESeq2 package, blind to design of the experiment. Top 10,000 most variable tag cluster were used to create the PCA plot. For PCA of individual CTSSs, top 8,000,000 most variable CTSSs were used.

### Heatmap visualisations

Heatmaps Bioconductor package (Perry M (18). heatmaps: Flexible Heatmaps for Functional Genomics and Sequence Features. R package version 1.2.0) was used to visualize motif locations or dinucleotide patterns across sequences. Sequences were in most cases centred on the SLIC-CAGE-identified dominant TSSs and ordered by the interquantile width of the corresponding tag cluster, placing the narrowest on top and the broadest tag cluster on the bottom of the heatmap. In the case of shifting promoters, the ordering was based on the distances between the dominant TSSs. Raw data with the dinucleotide matches was smoothed prior to plotting using the smoothing within the heatmaps package (Extended Data Figure 5a). Coverage plots (nucleosome positioning H3K4me3 ChIP-seq signal, CTSS or tag cluster coverage), were created using the CoverageHeatmap function from the Heatmaps package. The signal was weighted using the corresponding score (either nucleosome positioning score or CTSS/tag cluster log_10_(tpm + 1) value, as stated in the results and/or figure legends).

H3K4me3 ChIP-seq data that was used for gonadal germ cells E17.5M is under GEO Accession number GSM3426393 (SRR8029752 and SRR8029753)^44^. Biological replicates were merged prior to visualisation. H3K4me3 ChIP-seq data for MII oocyte is under GEO Accession number GSM2082662 (SRR3208745, SRR3208746, SRR3208747, SRR3208748)^45^. Biological replicates were merged prior to visualisation.

### Gene ontology (GO) enrichment analysis

Gene ontology enrichment analysis was performed using an over-representation test with clusterProfiler v.3.14.3^46^. Enriched GO terms associated with biological processes were determined based on a BH-adjusted *p*-value cutoff = 0.05. All annotated genes from TxDb.Mmusculus.UCSC.mm10.knownGene served as background

### Identification of TSS shifting promoters

Shifting promoters were identified for each sample in comparison to mESC cell line unless specified otherwise. The function shifting.promoters from the CAGEr package^26^ was used to identify shifting promoters, with a 3 tpm expression level and 0.01 false discovery rate (FDR) threshold, as previously described^4^.

### TATA-box and W-box motif analysis

TATA-box pwm from SeqPattern Bioconductor package was used in all analyses. Briefly, sequences centred on the dominant TSS were scanned with the TATA-box pwm. For TATA-box percentile match distribution plots, regions 35 to 21 bp upstream of the dominant TSSs were selected (15 bp), scanned with the TATA-box pwm in a sliding window and the highest pwm match percentile selected for each sequence.

Motif representations (seqlogo) were produced using the ggseqlogo package (CRAN), from sequences spanning 35 bp upstream and 5 bp downstream of the dominant TSSs. Lengths of W-box stretches were analysed using the seqinr (CRAN) package. Sequences spanning 34 to 23 bp upstream of the dominant TSSs were selected (12 bp), WW-box stretches identified in a 1 bp sliding window and the longest WW-box stretch selected per sequence. Reported are the distributions of longest W-box stretches per each sequence.

Tetranucleotide composition of TATA-like elements in shifting promoters was analysed in regions 34 to 23 bp upstream of the dominant TSSs. These regions were first scanned with a TATA-box pwm, and the highest 8 bp matching sequence (8 bp correspond to the width of the TATA-box pwm) within the 12 bp scanned region was selected for tetranucleotide composition analysis. Tetranucleotides were counted in a 1 bp sliding window. Presented are the top 10 tetranucleotides.

### Promoter GC-content calculation

GC-content of promoter regions in zebrafish was calculated using zebrafish developmental CAGE datasets mapped to danRer7 – egg and prim6 (ZebrafishDevelopmentalCAGE, R data package, data published in^47^). Samples were processed using CAGEr (as described above), and normalised consensus clusters extracted. Consensus clusters were resized to 2kb width using the consensus cluster centre as a fixing point, and GC-content calculated using the GC function (seqinr R package) on the extracted sequence. Analogously, GC-content was calculated for mouse promoter regions using consensus clusters generated using all SLIC-CAGE datasets.

### Expression profiling/classification using self-organising maps (SOMs)

SOM expression profiling was performed using kohonen R package for self-organising maps^48^. Only tag clusters located in promoter regions were used, i.e. tag clusters within 3kb of the annotated TSSs or in 5’UTR regions, and having a minimum of 1 tpm expression value in at least one sample (annotations as described TxDb.Mmusculus.UCSC.mm10.knownGene). Prior to clustering, the matrix of tag cluster expression values was scaled across samples without centring, and final SOM clusters were visualised using ggplot2.

Similarly, SOM clustering of dominant CTSSs was performed. Dominant CTSSs with at least 1 tpm expression value in at least one sample were selected prior to scaling and clustering.

### Nucleosome-preceding sequence motif analysis

To build the nucleosome-preceding position weight matrix we used mESC nAnT-iCAGE identified promoters. Nucleosome-preceding position weight matrix (pwm) was chosen as the most informative sequence part for nucleosome positioning as these sequences had the highest information content. Out of 20772 mESC promoters, we excluded 829 E16.5F vs mESC shifting promoters. We further divided the leftover mESC promoters into those with narrow (IQ-width <= 9, 7205 promoters) and broad transcription initiation (IQ-width >9, 12157). We further selected 10000 broad promoters to create the nucleosome-preceding pwm. Nucleosome-preceding sequences encompassed 6-50 bp downstream of the dominant TSSs. Out of leftover 2157 mESC broad promoters, we selected 829 to match the number of E16.5F vs mESC shifting promoters and serve as a test group in Extended Data Figure 12d. Sequences were further scanned with the nucleosome-preceding pwm using the seqPattern Bioconductor package, and the match scores visualised using the heatmaps Bioconductor package. Prior to producing metaplot, the match scores were winsorized to exclude extreme values (5^th^ to 95^th^ percentile of scores were selected). Relative scores for metaplots were then calculated by summing scores at each bp position in a stack of aligned sequences and dividing the sum with the number of sequences. Relatives scores were visualised using ggplot2.

## Data Availability

The data generated in this study has been submitted to ArrayExpress under accession number E-MTAB-9757 (https://www.ebi.ac.uk/arrayexpress/). Trackhub to visualise processed data in the UCSC genome browser will be made available.

## Code Availability

Custom analysis scripts are currently available at request and will be made publicly available on GitHub.

## Acknowledgments

This work was supported by The Wellcome Trust grant (106954) awarded to B.L. and F.M., MRC Core Funding (MC-A652-5QA10) and BBSRC Responsive Mode Grant (BB/R002703/1) awarded to H.G.L. N.C. was supported by an EMBO Advanced Long Term Fellowship (EMBO aALTF 626-2018) and a Wellcome Trust Seed Award (217309/Z/19/Z). B.L. is supported by the Medical Research Council UK (MC UP 1102/1). H.G.L acknowledges support from the National Institute for Health Research (NIHR) Imperial Biomedical Research Centre (BRC). Work in the Hajkova lab is supported by MRC funding (MC_US_A652_5PY70) and by an ERC grant (ERC-CoG-648879 – dynamic modifications). Y.H. is supported by the Japanese Society for the Promotion of Science (JSPS) Postdoctoral Fellowship (PD) No. 201700058. This study was also supported by the European Research Council (ERC) Advanced grant (ERC-2013-340551, Birtoaction) (to LT) and grant ANR-10-LABX-0030-INRT and a French State fund managed by the Agence Nationale de la Recherche under the frame program Investissements d’Avenir ANR-10-IDEX-0002-02 (to IGBMC). We also thank Laurence Game and the MRC LMS sequencing facility for support. Lastly, we thank Vedran Franke, Peter Sarkies and members of the Lenhard group for critical reading of the manuscript.

## Author contributions

N.C., B.L. and P.H designed the study. N.C. performed SLIC-CAGE and computational analyses. M.B., Y.H., H.G.L., P.H., L.T., S.V. and C.Y. provided PGC, GGC, oocyte and embryo samples. N.C., B.L. and P.H. wrote the manuscript with input by F.M. and all other authors.

## Materials & Correspondence

Correspondence and material requests should be to Nevena Cvetesic, Boris Lenhard or Petra Hajkova.

