## Extended Data for "Global regulatory transitions at core promoters demarcate the mammalian germline cycle"

### Extended Data Figures and Figure Legends

#### Extended Data Figure 1.

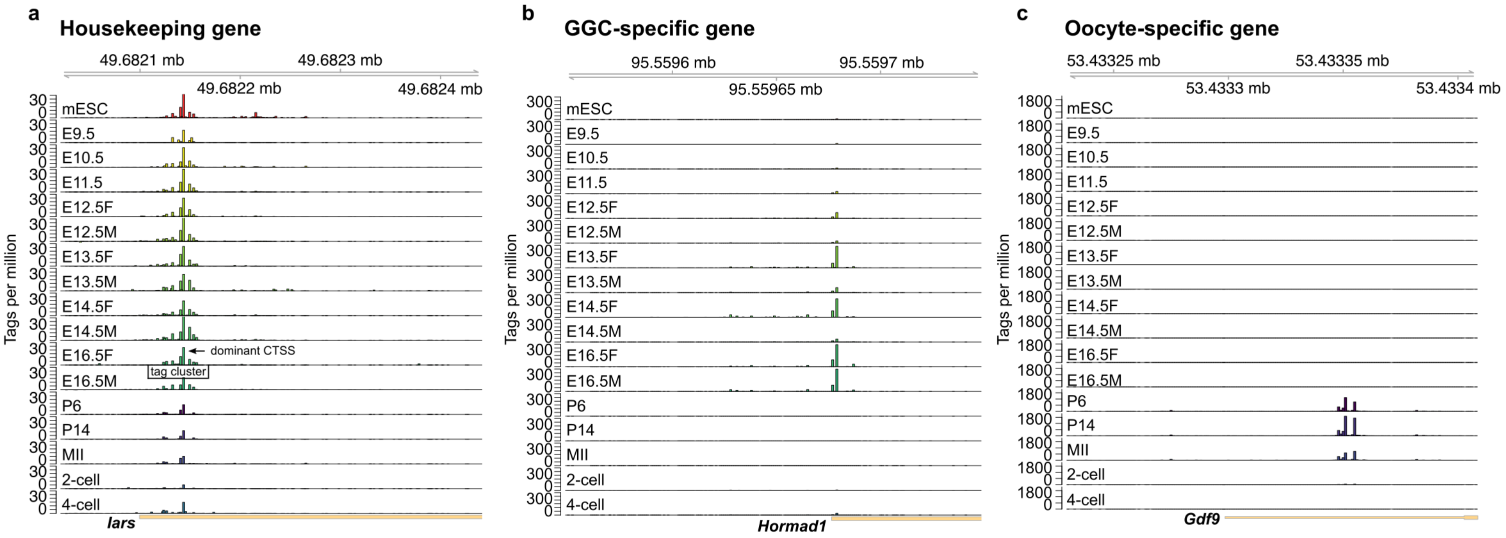


**Extended Data Figure 1. Uncovering dynamic gene expression at TSS resolution using SLIC-CAGE. a)** CTSS signal of an example housekeeping gene – isoleucyl-tRNA synthetase (*Iars*) Each bar represents an individual CTSS, and its height reflects the expression level. Cluster of CTSSs (tag cluster) and the most expressed CTSS in a tag cluster are labelled on the E16.5F sample. b) CTSS signal of an example gene expressed only in gonadal germ cells – Hormad Domain-Containing Protein 1 (*Hormad1*). c) CTSSs of an example gene expressed only in the oocytes – Growth Differentiation Factor (*Gdf9*). The colour scheme used in a) is used throughout the manuscript to represent the stages. Using SLIC-CAGE, we can identify individual CTSSs at 1 bp resolution and monitor gene expression dynamics.

**Extended Data Figure 2.**

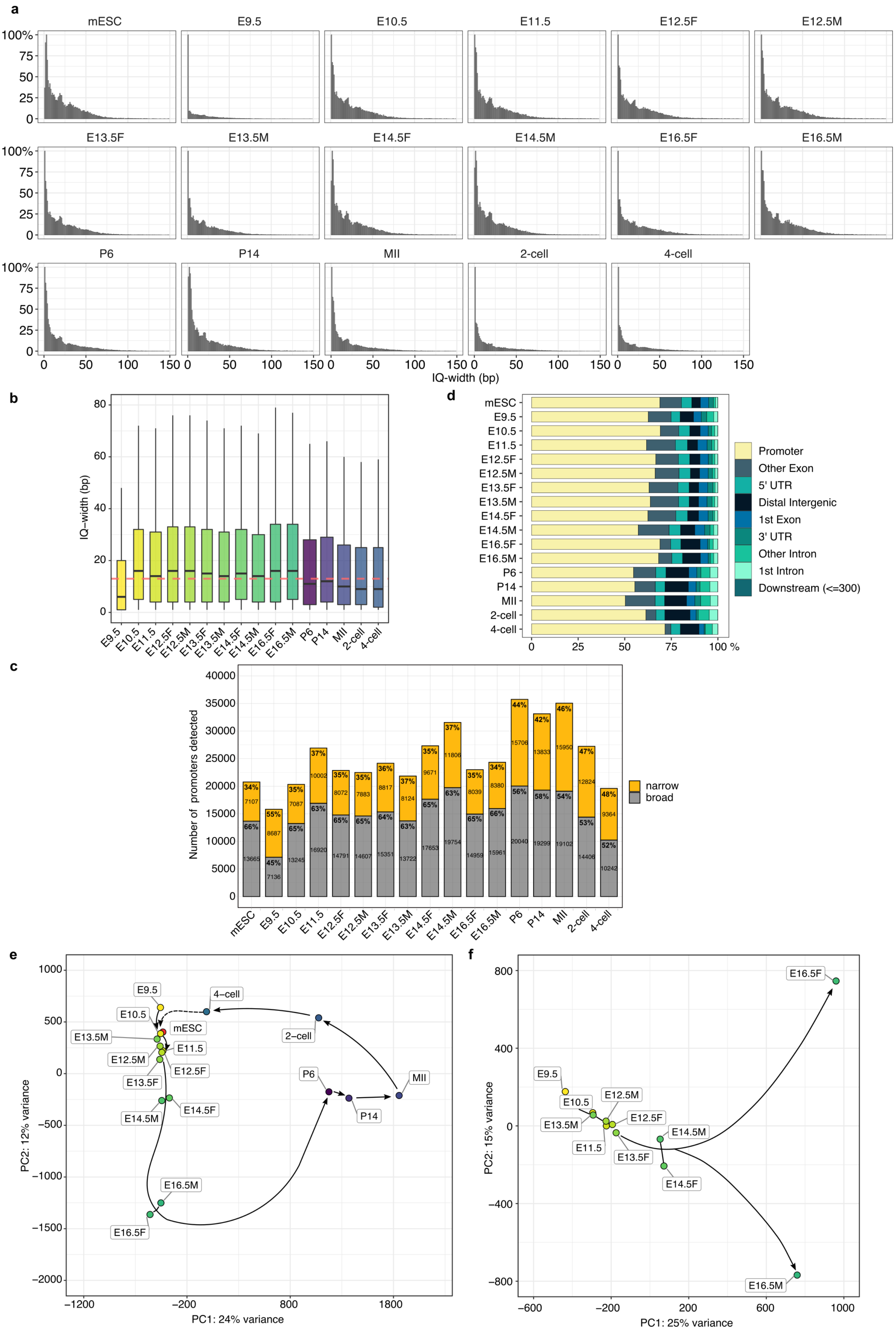

a
mESC
E9.5
E10.5
E11.5
E12.5F
E12.5M
E13.5F
E13.5M
E14.5F
E14.5M
E16.5F
E16.5M
P6
P14
MII
2-cell
4-cell
100%
75
50
25
0
0
50
100
150
IQ-width (bp)
b
IQ-width (bp)
80
60
40
20
0
E9.5
E10.5
E11.5
E12.5F
E12.5M
E13.5F
E13.5M
E14.5F
E14.5M
E16.5F
E16.5M
P6
P14
MII
2-cell
4-cell
d
mESC
E9.5
E10.5
E11.5
E12.5F
E12.5M
E13.5F
E13.5M
E14.5F
E14.5M
E16.5F
E16.5M
P6
P14
MII
2-cell
4-cell
0
25
50
75
100 %
Promoter
Other Exon
5' UTR
Distal Intergenic
1st Exon
3' UTR
Other Intron
1st Intron
Downstream (<=300)
c
Number of promoters detected
40000
35000
30000
25000
20000
15000
10000
5000
0
mESC
E9.5
E10.5
E11.5
E12.5F
E12.5M
E13.5F
E13.5M
E14.5F
E14.5M
E16.5F
E16.5M
P6
P14
MII
2-cell
4-cell
narrow
broad
e
PC2: 12% variance
PC1: 24% variance
1000
500
0
−500
−1000
−1500
−2000
−1200
−200
800
1800
E9.5
E10.5
mESC
E13.5M
E11.5
E12.5M
E12.5F
E13.5F
E14.5M
E14.5F
E16.5M
E16.5F
4−cell
2−cell
P6
P14
MII
f
PC2: 15% variance
PC1: 25% variance
800
400
0
−400
−800
−600
−200
200
600
1000
E16.5F
E9.5
E10.5
E12.5M
E12.5F
E13.5M
E11.5
E13.5F
E14.5M
E14.5F
E16.5M

**Extended Data Figure 2. Transcription initiation at 1 bp resolution in mouse PGCs, GGCs, oocytes and embryos. a)** Distribution of tag cluster IQ-widths (width of the region encompassing central 80% of the signal). All tag clusters identified from merged biological replicates were used. Narrow tag clusters in E9.5 PGCs imply a lower complexity of the library, as the starting total RNA amount was below SLIC-CAGE sensitivity. **b)** Boxplot representation of IQ-widths from a). **c)** Number and percentage of narrow (IQ-width < 9 bp) and broad promoters (IQ-width >= 9 bp) detected in each sample. **d)** Genomic locations of SLIC-CAGE identified tag clusters. **e)** PCA of all stages based on expression of all identified CTSSs. **f)** PCA separation of PGCs and GGCs based on expression of all identified CTSSs, showing good separation of the late gonadal germ cell stages. Black lines and arrows mark the differentiation time-course.

**Extended Data Figure 3.**

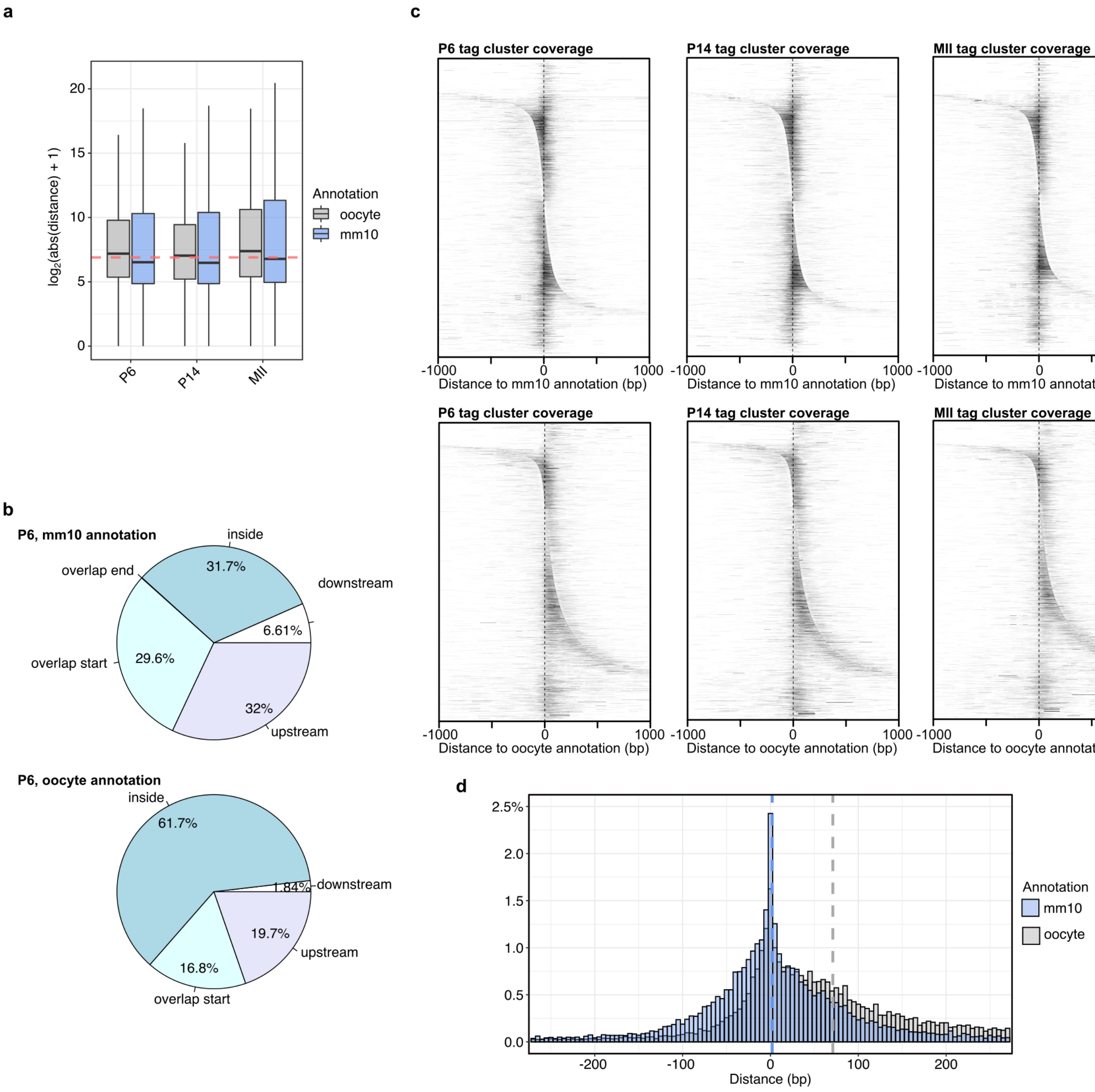


**Extended data Figure 3. Comparison of SLIC-CAGE identified CTSS locations with the mm10-based and oocyte transcript annotation** [1]. **a)** Distribution of absolute distance of SLIC-CAGE-identified tag clusters in P6, P14 or MII oocytes, from the TSSs defined by the mm10 or Veselovska et al oocyte transcript annotation ($log_2$ scale). The horizontal red dashed line is the median. **b)** Genomic locations of SLIC-CAGE tag clusters. Locations are defined by mm10 or Veselovska et al oocyte transcript annotation. **c)** Heatmap visualisation of SLIC-

CAGE-identified tag clusters, centred on the mm10 TSSs (top) or Veselovska oocyte TSSs (bottom) and sorted by the distance from the mm10 or Veselovska TSSs. The scale is the same in all heatmaps and reflects the P6 tag cluster expression ($\log_2$(tpm + 1)). **d)** Distribution of SLIC-CAGE-identified P6 tag cluster distance upstream (negative distance) or downstream (positive distance) from mm10 or oocyte annotation defined TSSs. Shorter average distance from the mm10 annotation (a, d), higher percentage overlap with the annotated promoter starts (b) and symmetrical distances from the annotated TSSs (c, d) show that SLIC-CAGE-identified tag clusters are closer to the mm10 annotation. This substantiates the need for a new accurate oocyte transcript annotation based on the SLIC-CAGE identified TSSs.

**Extended Data Figure 4.**

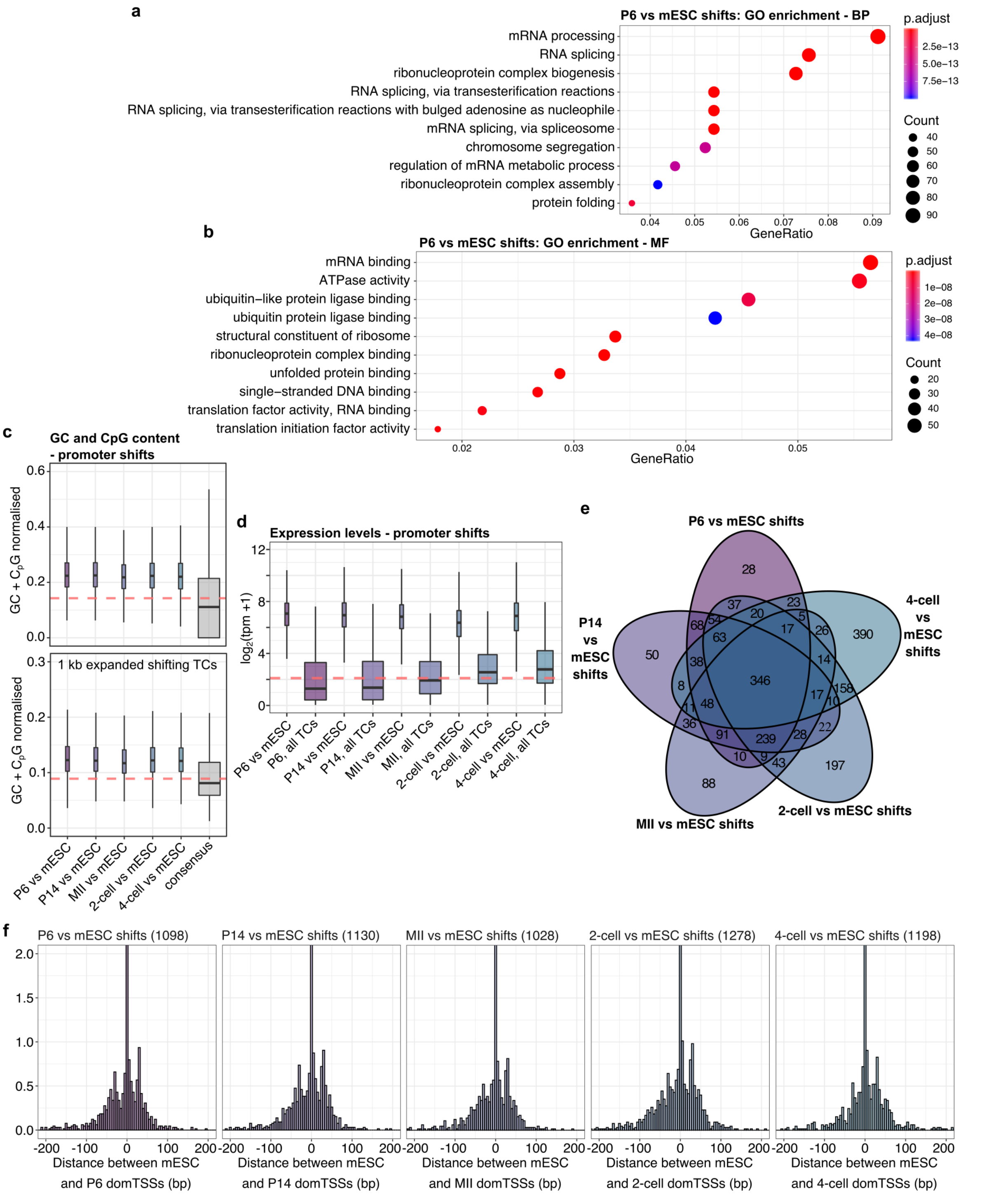


**Extended Data Figure 4. Identification of oocyte and early embryo shifting promoters.** GO enrichment of P6 vs mESC shifting promoters: **a)** biological process, **b)** molecular function. These results demonstrate that P6 vs mESC shifting promoters are largely housekeeping, i.e. ubiquitously expressed genes. **c)** Distribution of GC + CpG content of SLIC-

CAGE identified tag clusters. Top panel - number of GC + CpG dinucleotides in identified tag clusters, normalised by tag cluster width. Bottom panel – number of GC + CpG dinucleotides in tag clusters flanked with additional 1 kbp (500 bp upstream and downstream), normalised by the width of the expanded regions. Shifting promoters have a higher GC + CpG content. The horizontal red dashed line is the median. **d)** Expression in shifting or all identified promoters (marked with all TCs). Shifting promoters are higher expressed than the average, as expected for ubiquitously expressed CG-rich promoters. **e)** Overlap of shifting promoters identified through pairwise comparison of CTSSs from each stage and mESC. Large overlap of oocyte vs mESC and early 2-cell embryo vs mESC shifts is due to the maternally inherited RNA in the early 2-cell embryo (pre-major ZGA). **f)** Distribution of distances in shifting promoters between mESC and P6, P14 oocyte or early 2-cell or 4-cell embryo dominant TSSs. Shifts occur both upstream and downstream of the mESC-identified dominant TSSs, with a slight upstream preference.

**Extended Data Figure 5.**

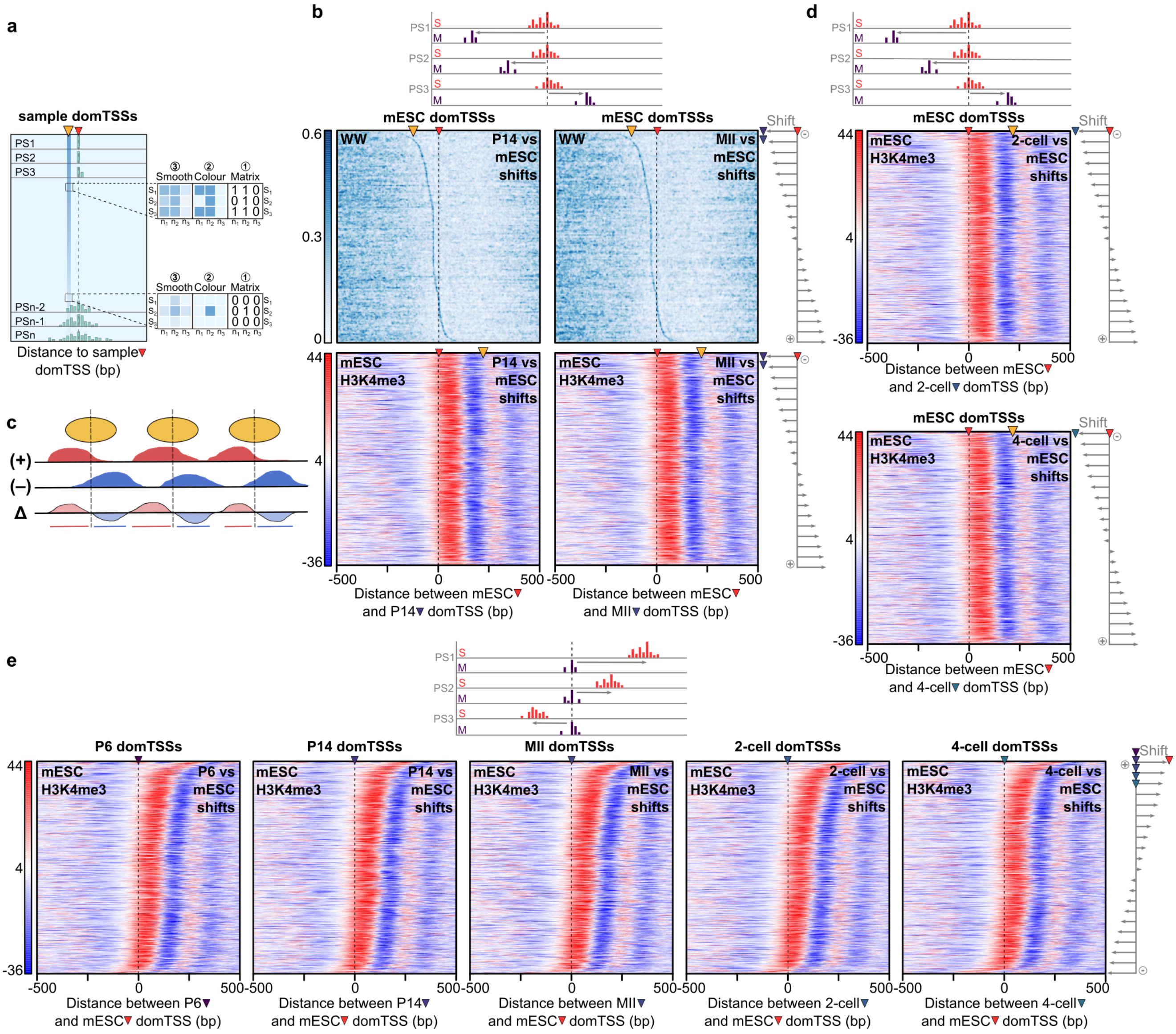


**Extended Data Figure 5. Somatic-maternal-somatic transitions of transcription initiation in shifting promoters a)** Scheme of a heatmap showing WW dinucleotide density in promoter sequences. The sequences are centred on the dominant CTSSs and sorted according to the IQ-width (sharp promoters are on top of the heatmap, and broad are on the bottom; green bars within the heatmap represent individual CTSSs; position of the dominant TSSs is marked with a red arrow; throughout the manuscript, the colour of the arrow corresponds to the sample colour from Extended Data Figure 1a). PS1-PSn denotes individual promoter sequences, while the orange arrowhead indicates WW enrichment at the expected position of the TATA-box/TATA-like element (30 bp upstream of the dominant TSS). Throughout the manuscript, orange arrowhead is used to emphasize a signal or a region. The insets show calculation and visualisation of the dinucleotide patterns. Genomic sequences (S1-S3) are aligned and sorted in a matrix-type representation (marked with 1). Presence of a WW dinucleotide at a certain position (n1-n3) is marked with 1 and the absence with 0. The binary matrix (labelled with 1) is not directly visualized (as labelled with 2); instead, 2D binned kernel density estimate is applied to the matrix and the new values are then mapped to different shades of blue (labelled with 3, smooth). **b)** WW density (top) and mESC H3K4me3 signal coverage (bottom) from reads mapping to the plus (+) and minus (-) strand (schematics explaining the data is in c) of P14 or MII vs mESC shifting promoters, centred to the mESC dominant TSSs (marked with a red arrow) and sorted by the distance and orientation of the shift (scheme on the top and right). Orange arrowheads in the WW heatmaps indicate the WW enrichment, and in the H3K4me3 heatmaps the internucleosomal region. WW enrichment follows P14 and MII dominant TSSs, while the dominant TSS positions in mESC are aligned to the H3K4me3 marked +1 nucleosome. **c)** Schematics of the subtracted H3K4me3 coverage (Δ) of reads mapping to the plus (+) and minus (–) strand, demonstrating how the signal corresponds to +1 nucleosome positioning. **d)** Subtracted mESC H3K4me3 signal visualised as coverage in the early 2-cell and 4-cell vs mESC shifting promoters. Heatmaps are centred on the mESC dominant CTSSs and sorted by the distance and orientation of the shift (schemes on top and right). mESC dominant TSSs, and not 2-cell or 4-cell, are aligned to the +1 nucleosomes marked with H3K4me3. **e)** Subtracted mESC H3K4me3 coverage visualised in P6, P14, MII, early 2-cell and 4-cell vs mESC shifting promoters centred to sample-identified dominant TSSs and sorted by the distance and orientation of the shift (schemes on top and right). This again shows that mESC H3K4me3 marked +1 nucleosomes direct mESC dominant TSS positions.

**Extended Data Figure 6.**

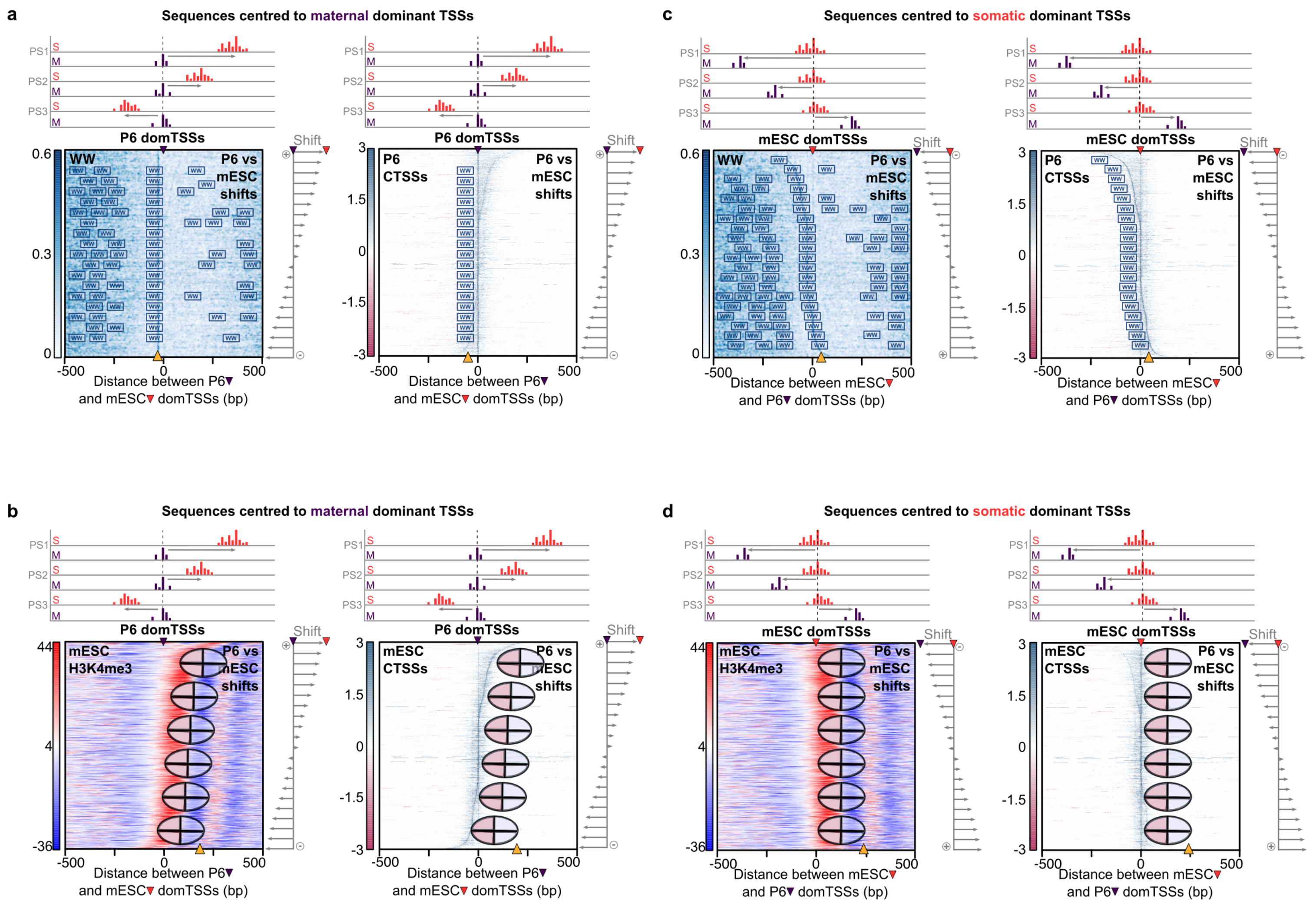


**Extended Data Figure 6. Schemes explaining TSS shifts, reflecting somatic-maternal-somatic grammar transitions.** Heatmaps are the same as presented in Figure 2 and Extended Data Figure 5, albeit with schematics overlaid. All sequences are ordered by the distance and orientation of the shift (schemes on top and right of heatmaps). **a)** WW density (left) and P6 CTSS coverage (right) in P6 vs mESC shifts centred on the P6 dominant TSSs (marked with a purple arrowhead on top of heatmaps). Scale in WW heatmaps represents the WW density and in CTSS heatmaps represents expression ($\log_2$(tpm +1)) - sense CTSS signal is shown in blue (positive) and antisense in red (negative). Schematic WW-boxes overlay the WW heatmap, and the WW-boxes directing transcription initiation are marked with an orange arrowhead at the bottom. The CTSS heatmap (right) shows that the P6 dominant (strongest) CTSSs are aligned with the WW-boxes 30 bp upstream. **b)** mESC H3K4me3 (left) and mESC CTSS coverage in P6 vs mESC shifts centred on the P6 dominant TSSs. Schematic +1 nucleosomes are overlaid with the H3K4me3 coverage signal and the internucleosomal region marked with an orange arrowhead on the bottom of the heatmap (see Extended Data Figure 5c for scheme of the H3K4me3 signal). mESC CTSS heatmap (right) demonstrates that the mESC dominant CTSSs are aligned with the +1 nucleosomes **c)** Same as in a) but centred on the mESC dominant CTSSs, demonstrating again that the P6 dominant CTSSs in P6 vs mESC shifting promoters align with the W-boxes. d) Same as in b), but centred on the mESC dominant CTSS, demonstrating again that the mESC dominant CTSSs in P6 vs mESC shifts align with the +1 nucleosomes.

**Extended Data Figure 7.**

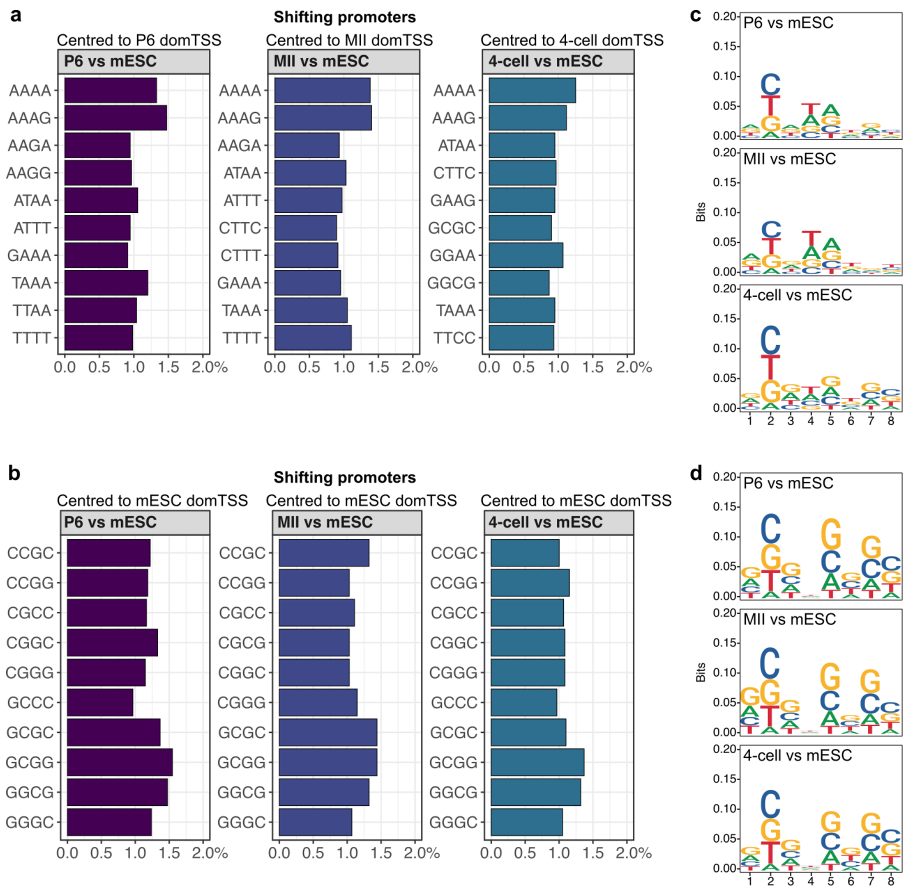


**Extended Data Figure 7. Sequence analysis of TATA-like elements in shifting promoters.** Regions 34 to 23 bp upstream of the dominant TSSs were scanned with a TATA-box pwm in P6, MII oocyte and 4-cell embryo vs mESC shifting promoters. The highest matching sequence (8 bp) within the scanned region (12 bp width) was selected and its tetranucleotide composition analysed. Presented are top 10 tetranucleotides in shifting promoters uncovered when sequences are centred on **a)** P6, MII oocyte or 4-cell embryo dominant TSSs; **b)** mESC dominant TSSs. Tetranucleotide composition upstream of the P6 or MII dominant TSSs (a) is more AT-rich than upstream of the mESC dominant TSSs in the same set of shifting promoters. This supports the notion that the W-box dictates the dominant TSS position in the P6 and MII oocyte. 4-cell vs mESC shifting promoters include leftover maternal transcripts, hence tetranucleotides reflect a combination of the AT-rich W-box and GC-rich nucleosome positioning signal. **c)** Sequence logo of top scoring TATA-box pwm matched sequences in P6, MII or 4-cell embryo vs mESC shifts centred on P6, MII or 4-cell embryo dominant TSSs (8 bp match in a 12 bp wide region). **d)** Sequence logos of top scoring TATA-box pwm matched sequences (presented in a) and b) )in P6, MII or 4-cell embryo vs mESC shifts, centred on **c)** P6 dominant TSSs; **d)** mESC dominant TSSs.

### Extended Data Figure 8

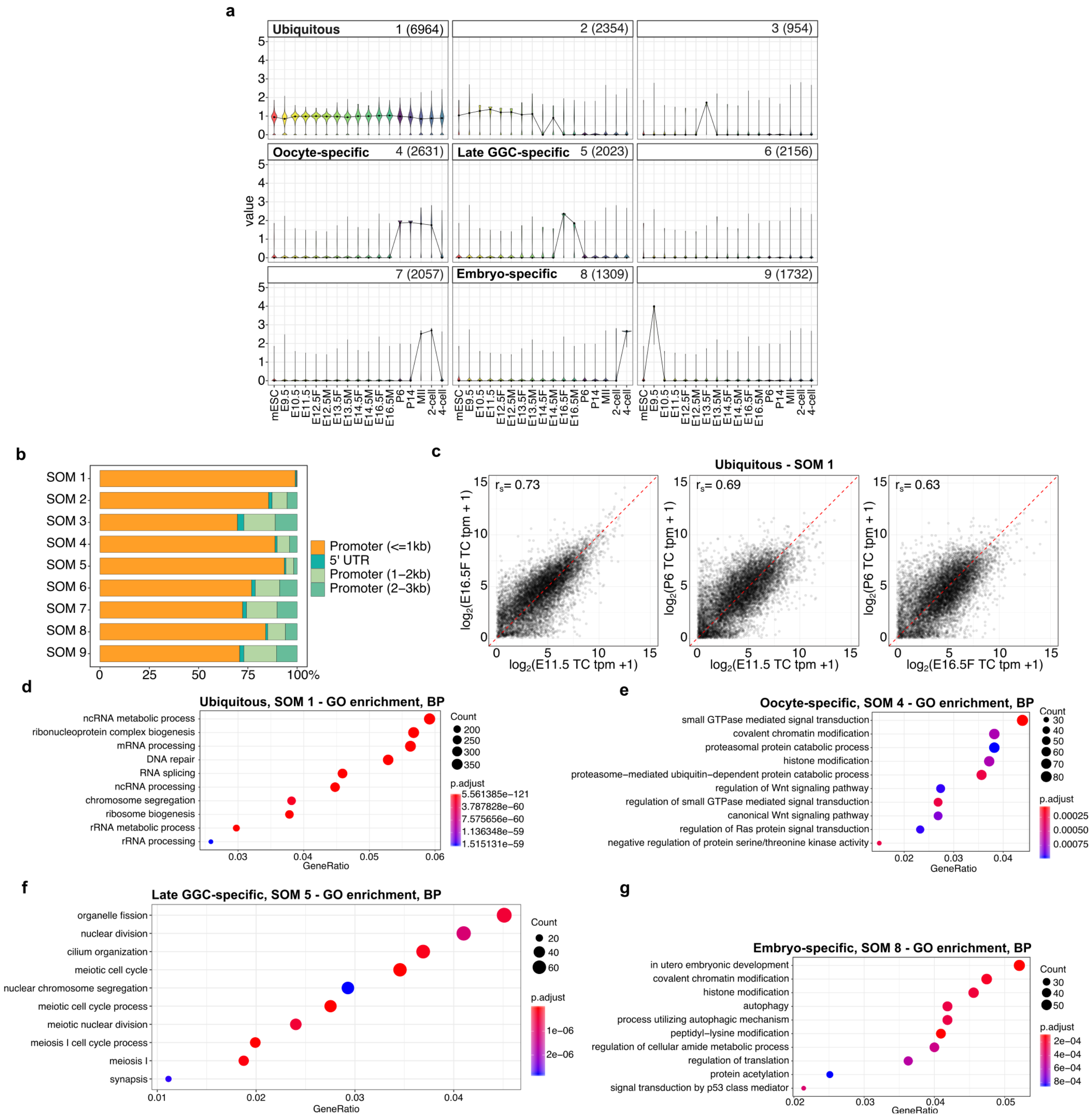


**Extended Data Figure 8. SOM clustering of SLIC-CAGE-identified promoters. a)** SOM-identified promoter clusters. Ubiquitously expressed, oocyte-specific, late-gonadal germ cell-specific and embryo-specific promoters are highlighted. **b)** Genomic locations of tag clusters used for SOM clustering (defined by the distance from the annotated TSSs; only promoter related tag clusters were used for SOM). **c)** Expression correlation of ubiquitous tag clusters

(SOM 1) in example samples (GGCs E11.5, E16.5F and oocyte P6) demonstrating that ubiquitous genes retain similar expression levels across stages regardless of grammar switch. Spearman correlation coefficient is shown in top left corner. **d-g)** GO enrichment (biological process terms) of the ubiquitously expressed SOM cluster 1 (d), oocyte specific SOM cluster 4 (e), late gonadal germ cell-specific SOM cluster 5 (f) and embryo specific SOM cluster 8 (g). Biological process terms confirm the biological relevance of promoter-level SOM clustering into – ubiquitous, oocyte-specific, late GGC-specific and embryo-specific.

**Extended Data Figure 9.**

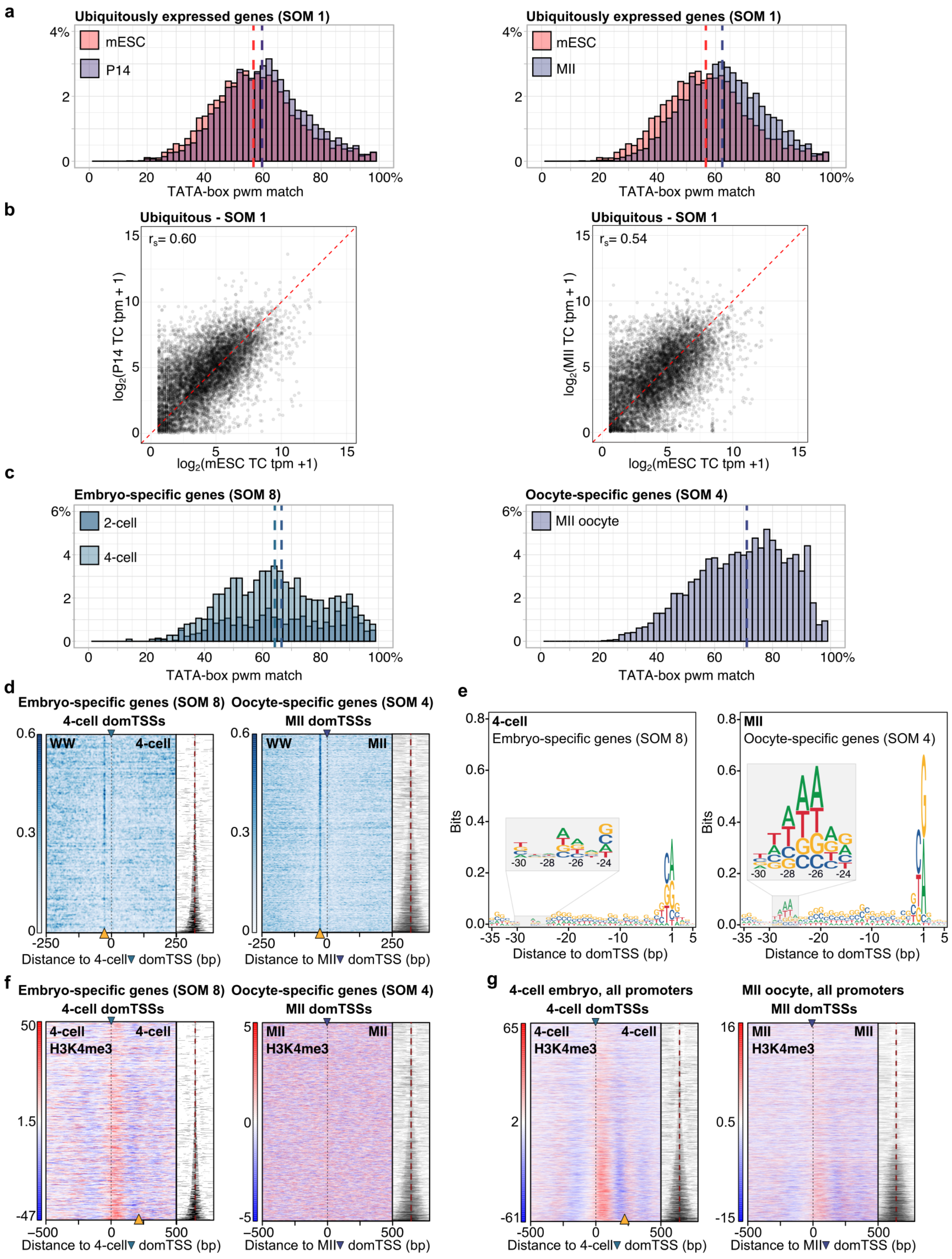


**Extended Data Figure 9. Sequence features of embryo- and oocyte-specific genes. a)** Distributions of the TATA-box pwm percentile matches in ubiquitously expressed promoters (SOM 1, Extended Data Figure 8a) centred on the P14- , MII or mESC-identified dominant TSSs. Percentile matches are calculated in a 1 bp sliding window spanning 35 to 21 bp upstream of the dominant TSSs. Best match is reported per each sequence. **b)** Expression correlation of ubiquitously expressed tag clusters (SOM 1) in mESC and P14 oocyte, or mESC and MII oocyte. Spearman correlation coefficient is reported. This demonstrates that ubiquitously expressed genes retain similar expression levels across differentiation stages. **c)** Distributions of the TATA-box pwm percentile matches in the embryo or oocyte specific genes (centred on 2-cell or 4-cell embryo dominant TSSs – left, or MII oocyte dominant TSSs – right). Calculations are as in a) **d)** WW density in embryo-specific promoters (SOM 8, Extended Data Figure 8a) centred on the 4-cell embryo dominant TSSs (left), and oocyte-specific promoters (SOM cluster 4, Extended Data Figure 8a) centred on the MII dominant TSSs. Sequences are ordered by the IQ-width (right of heatmap, tag cluster coverage in grey visualises the IQ-width; the dashed red vertical line represents the dominant TSS position). The orange arrowheads mark the positions of the WW enrichment, 30 bp upstream of the dominant TSS. The arrowhead in a sample-specific colour on top of the heatmaps marks the dominant TSSs (colour-scheme from Extended Data Figure 1). **e)** Sequence logos produced from a region encompassing 35 bp up- and 5 bp downstream of the dominant TSSs of embryo- or oocyte-specific promoters. **f)** Heatmaps showing 4-cell embryo H3K4me3 or MII oocyte H3K4me3 coverage in embryo- (left) or oocyte-specific (right) promoters. Sequences are centred on the sample-specific dominant TSSs (marked with an arrowhead) and ordered by the IQ-width (right of heatmap). **g)** Same as e), however using all SLIC-CAGE-identified promoters.

**Extended Data Figure 10.**

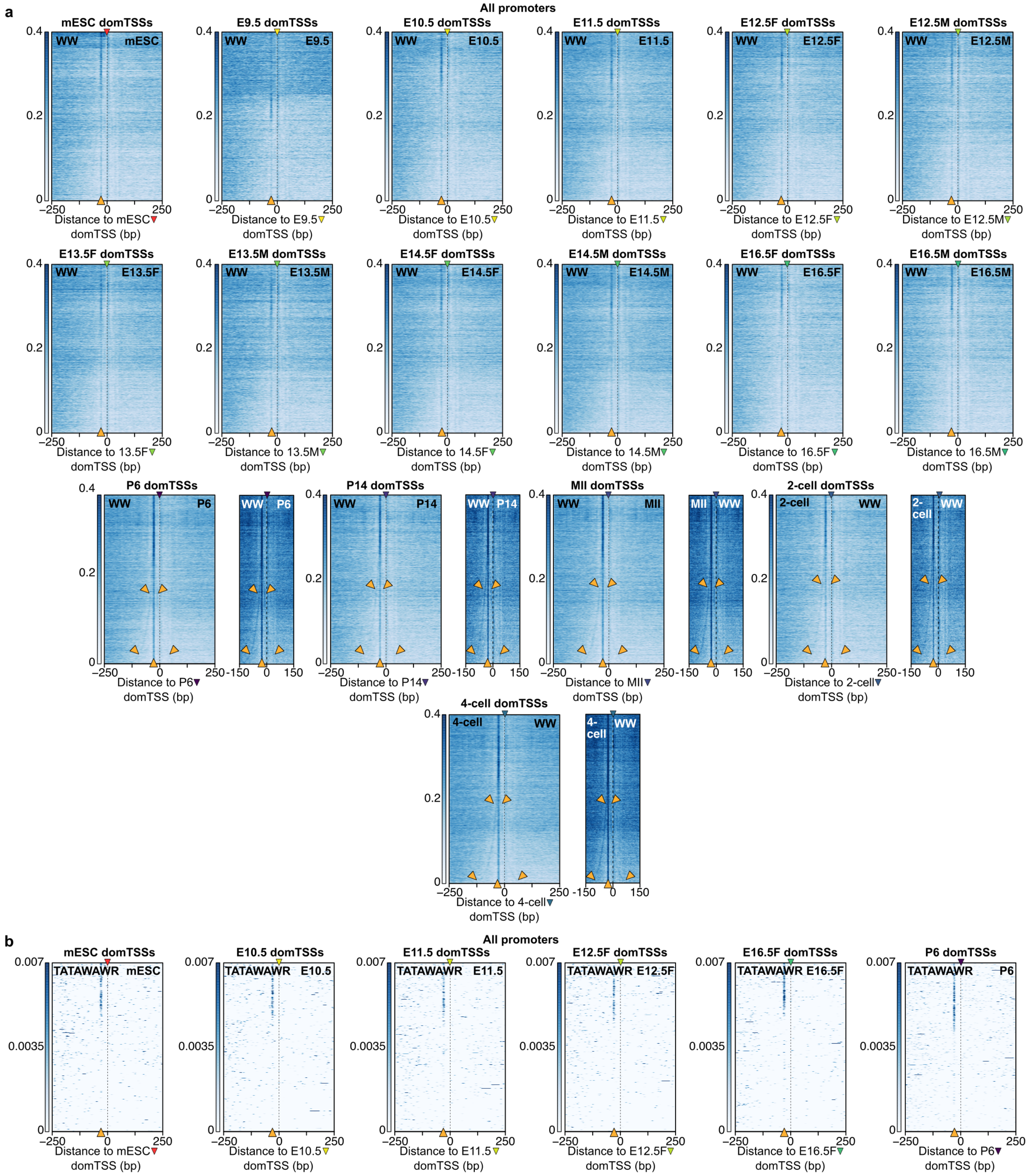


**Extended Data Figure 10. Sequence patterns in all SLIC-CAGE identified promoters.**

Sequences in a) and b) are centred on the sample-specific dominant TSSs (marked by an arrowhead on top of heatmap) and ordered by the tag cluster IQ-width, with narrower tag clusters at the top, and broader at the bottom of the heatmaps. **a)** WW density (top) in all SLIC-CAGE identified promoters. Orange arrowheads on the bottom of the heatmaps indicate the position of the WW enrichment. i.e. putative TATA-boxes/TATA-like elements. Additional orange arrowheads in the P6, P14, MII, 2-cell and 4-cell heatmaps indicate a weak diverging WW enrichment, likely representing the most outer W-box of the multiple W-box broad promoter architectures (see Main). To better visualise these diverging WW enrichments, these sample are in addition shown on a narrower WW heatmap encompassing a 300 bp region around the dominant TSS on a different scale (all 300 bp window heatmaps are on the same scale, and separately, all 500 bp heatmaps are on the same scale). **b)** Density of the consensus TATAWAWR TATA-box motif. The orange arrowhead at the bottom of the heatmap indicates the position of the putative motif.

**Extended Data Figure 11.**

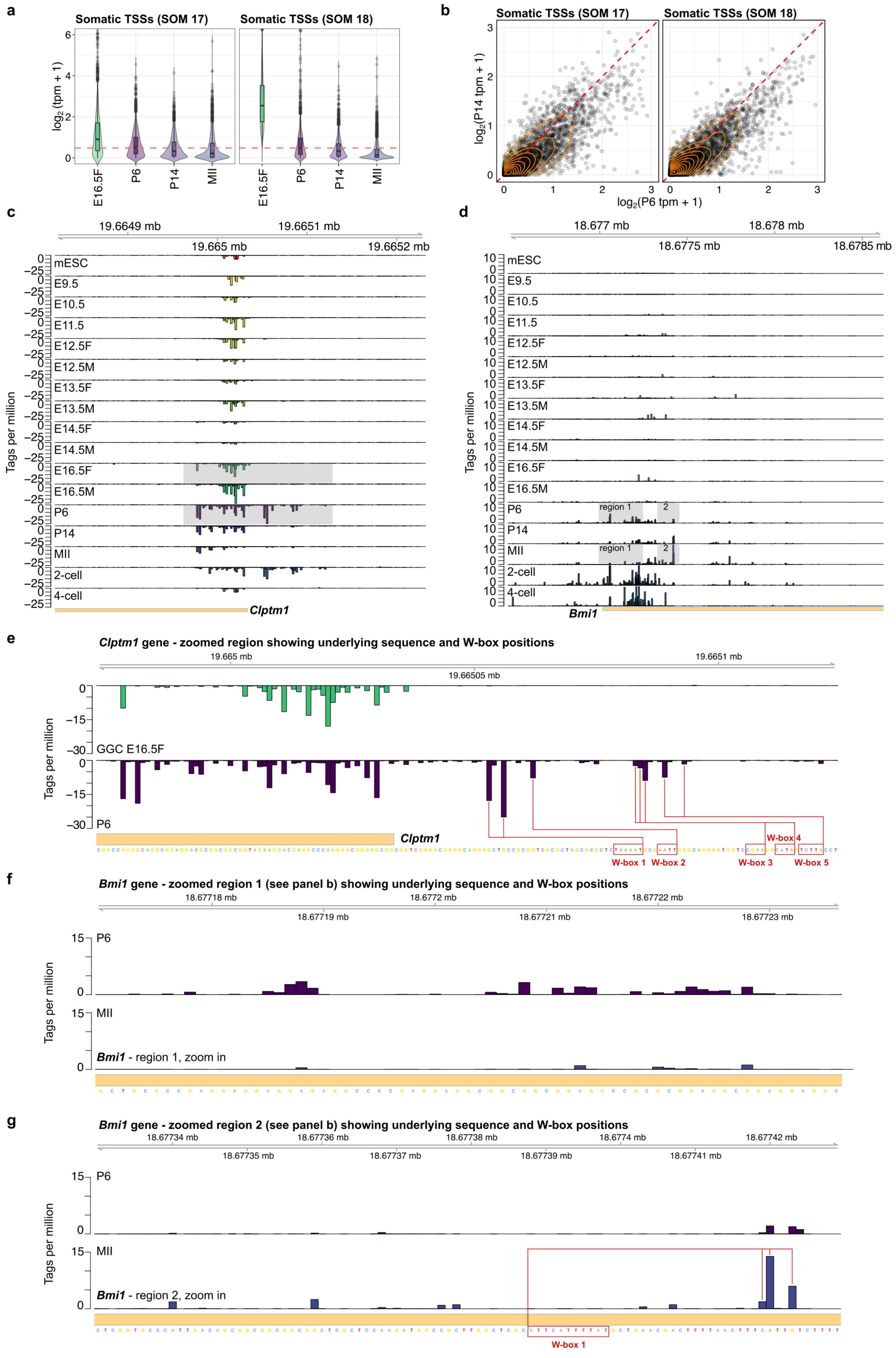

a
Somatic TSSs (SOM 17)
Somatic TSSs (SOM 18)
log2 (tpm + 1)
E16.5F
P6
P14
MII
b
log2(P14 tpm + 1)
log2(P6 tpm + 1)
c
19.6649 mb
19.6651 mb
19.665 mb
19.6652 mb
mESC
E9.5
E10.5
E11.5
E12.5F
E12.5M
E13.5F
E13.5M
E14.5F
E14.5M
E16.5F
E16.5M
P6
P14
MII
2-cell
4-cell
Tags per million
Clptm1
d
18.677 mb
18.678 mb
18.6775 mb
18.6785 mb
region 1
2
Bmi1
e
Clptm1 gene - zoomed region showing underlying sequence and W-box positions
19.665 mb
19.6651 mb
19.66505 mb
GGC E16.5F
W-box 1
W-box 2
W-box 3
W-box 4
W-box 5
f
Bmi1 gene - zoomed region 1 (see panel b) showing underlying sequence and W-box positions
18.67718 mb
18.6772 mb
18.67722 mb
18.67719 mb
18.67721 mb
18.67723 mb
Bmi1 - region 1, zoom in
g
Bmi1 gene - zoomed region 2 (see panel b) showing underlying sequence and W-box positions
18.67734 mb
18.67736 mb
18.67738 mb
18.6774 mb
18.67742 mb
18.67735 mb
18.67737 mb
18.67739 mb
18.67741 mb
Bmi1 - region 2, zoom in

**Extended Data Figure 11. Non-maternal transcription initiation in the P6 oocyte. a)** Expression of somatic dominant CTSSs in E16.5F GGCs and P6, P14 and MII oocytes (SOM clusters 17 and 18, see Extended Data Figure 16a). **b)** Scatter plot of somatic dominant CTSSs expressed in P6 versus P14 oocyte stage (SOM17 - left, and SOM18 – right). Orange lines display 2D density contours. **c, d)** CTSS signal of genes with non-maternal transcription initiation in the P6 oocyte. Each bar represents an individual CTSS, and its height reflects the expression level. **c)** Cleft lip and palate transmembrane protein 1 - *Clptm1*. CTSS signal in the P6 oocyte is reminiscent of the signal in the early 2-cell embryo where it likely represents a transition from the maternal to somatic grammar. Shaded region is presented with the underlying sequence in e). **d)** Polycomb complex protein BMI-1 – *Bmi1*. CTSS signal in the P6 oocyte is again similar to the signal in the early 2-cell embryo. Shaded regions 1 and 2 are presented with the underlying sequence in f) and g), respectively. **e)** Magnified view of the *Cpltm1* promoter region in the E16.5F GGCs and P6 oocyte. Sequence is shown below the CTSS signal and the putative W-boxes driving novel transcription initiation in the P6 oocyte are marked in red. No putative W-boxes were identified for the CTSSs similarly expressed in both E16.5F and P6 stages, indicating a non-maternal transcription initiation in the P6 oocyte. **f)** Magnified view of the *Bmi1* promoter region 1 (as highlighted in b) in the P6 and MII oocyte. No putative W-boxes were identified, indicating a non-maternal transcription initiation in the P6 oocyte. Accordingly, the CTSS signal at these locations is lower in the P14 and MII oocyte. **g)** Magnified view of the *Bmi1* promoter region 2 (as highlighted in b) in the P6 and MII oocyte. Sequence is shown below the CTSS signal and the putative W-boxes driving transcription initiation in the MII oocyte are marked in red.

**Extended Data Figure 12.**

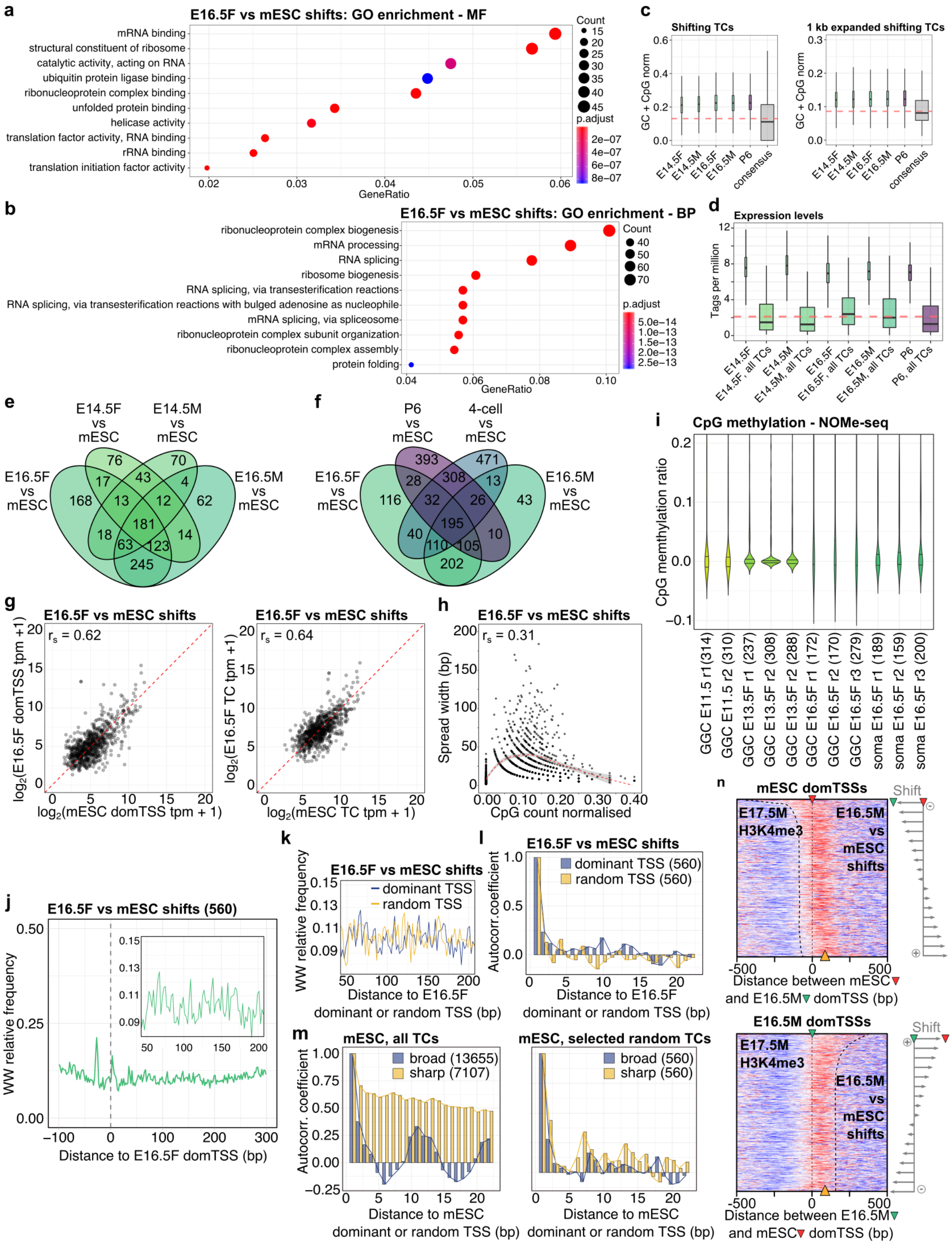


**Extended Data Figure 12. Sequence features underlying the GGC transcription initiation transitions.** GO enrichment analysis of the E16.5F vs mESC shifting promoters – **a)** molecular function, **b)** biological process. E16.5F vs mESC shifts are enriched in housekeeping terms. **c)** GC + CpG content of shifting promoters– number of GC + CpG is normalised by tag cluster width (left) or the width of tag clusters flanked with additional 500 bp upstream and downstream regions (right). Sample labels (below) represent shifting promoters identified between each sample and mESCs, while consensus refers to consensus promoters (based on all samples). Shifting promoters have a higher GC + CpG content than average. **d)** Expression of shifting or all identified promoters (marked with all TCs). E16.F shifting promoters are higher expressed than the average. **e)** Overlap of the GGC identified shifting promoters (E14.5F/M or E16.5F/M vs mESC). There is a large overlap of shifting promoters in the E16.5 and the preceding E14.5 stage, both male and female, demonstrating that: 1) the origin of late GGCs shifts precedes the E16.5 stage; 2) these shifts are not sex specific. **f)** Overlap of GGC, oocyte and 4-cell embryo vs mESC shifts. The majority of promoters that shift from the canonical somatic code in the E16.5F stage, also shift from the somatic to the maternal code (P6). The overlap of the 4-cell vs mESC and P6 vs mESC shifts is likely due to the leftover maternal RNA. **g)** Correlation of the CTSS (left) or tag cluster (right) expression levels in the GGC E16.5F vs mESC shifts. Shifting promoters generally retain a similar expression level prior to and after shifting to a different TSS pattern. **h)** Correlation of TSS spread width in the E16.5F vs mESC shifting promoters that became broader than in mESC, with the GC + CpG dinucleotide content within the spread region. Spearman correlation coefficient is shown. GC + CpG dinucleotide count is normalised using the spread width. Spread width of the CTSS signal slightly correlates with the GC + CpG content within the spread region. **i)** CpG methylation levels measured by NOMe-seq in the regions of TSS spread. As expected for active promoters, methylation level is low in GGC E16.5F spread regions in all samples. **j)** WW dinucleotide frequency in E16.5F vs mESC shifting promoters selected to include at least 1 bp distance between the dominant TSSs (560 E16.5F vs mESC shifts). Sequences are centred on the E16.5F dominant TSSs. The inset shows a magnified view of the region encompassing 50-200 bp downstream of the dominant TSSs. WW-periodicity indicative of the precisely positioned +1 nucleosome is weak due to a low number of sequences (see panel m). **k)** WW dinucleotide frequency in the E16.5F vs mESC shifting promoters selected as in j), centred on the true dominant CTSSs or a random CTSSs selected from the shifting promoters. WW dinucleotide periodicity, although overall weak, appears stronger in the E16.5F when centred on the true dominant than a random CTSSs, indicating a true biological signal for +1 nucleosome positioning. **l)** Autocorrelation analysis of the WW dinucleotide frequency in sequences 50-200 bp downstream of the dominant CTSSs in E16.5F vs mESC shifts selected as in j). Sequences were centred on the true dominant CTSSs (purple), or random CTSSs selected from the shifting promoter. Autocorrelation signal confirms the WW periodicity identified in j) – periodicity exhibits a maximum at about 10 bp, apparent only when centred on the true dominant CTSSs. **m)** Autocorrelation analysis of the WW frequency in: 1) all mESC promoters (left) separated into sharp (7107 promoters, IQ-width < 9bp) and broad (13655 promoters, IQ-width >= 9bp) and centred on the dominant CTSSs; 2) 560 randomly selected broad and sharp promoters (right). This control shows the autocorrelation signal for canonical broad promoters, where the precise +1 nucleosome positions determine the position of the dominant CTSSs. Subsampling the sequences to the same number as in panel l) significantly weakens the signal, similarly to the E16.5F shifting sequences in l). **n)** Heatmaps showing E17.5M H3K4me3 signal coverage in E16.5M vs mESC shifting promoters, centred on the mESC (top) or E16.5M dominant CTSSs (bottom) and ordered by the distance and orientation of the shift (scheme on the right). The orange arrowhead (bottom) marks the position of the +1 nucleosome, while the black dashed line contours its position. The H3K4me3 marked +1 nucleosome signal is weak, however it aligns well with both mESC and E16.5M dominant CTSSs, supporting that canonical and alternative +1 nucleosome positions guide transcription initiation in late GGCs.

### Extended Data Figure 13.

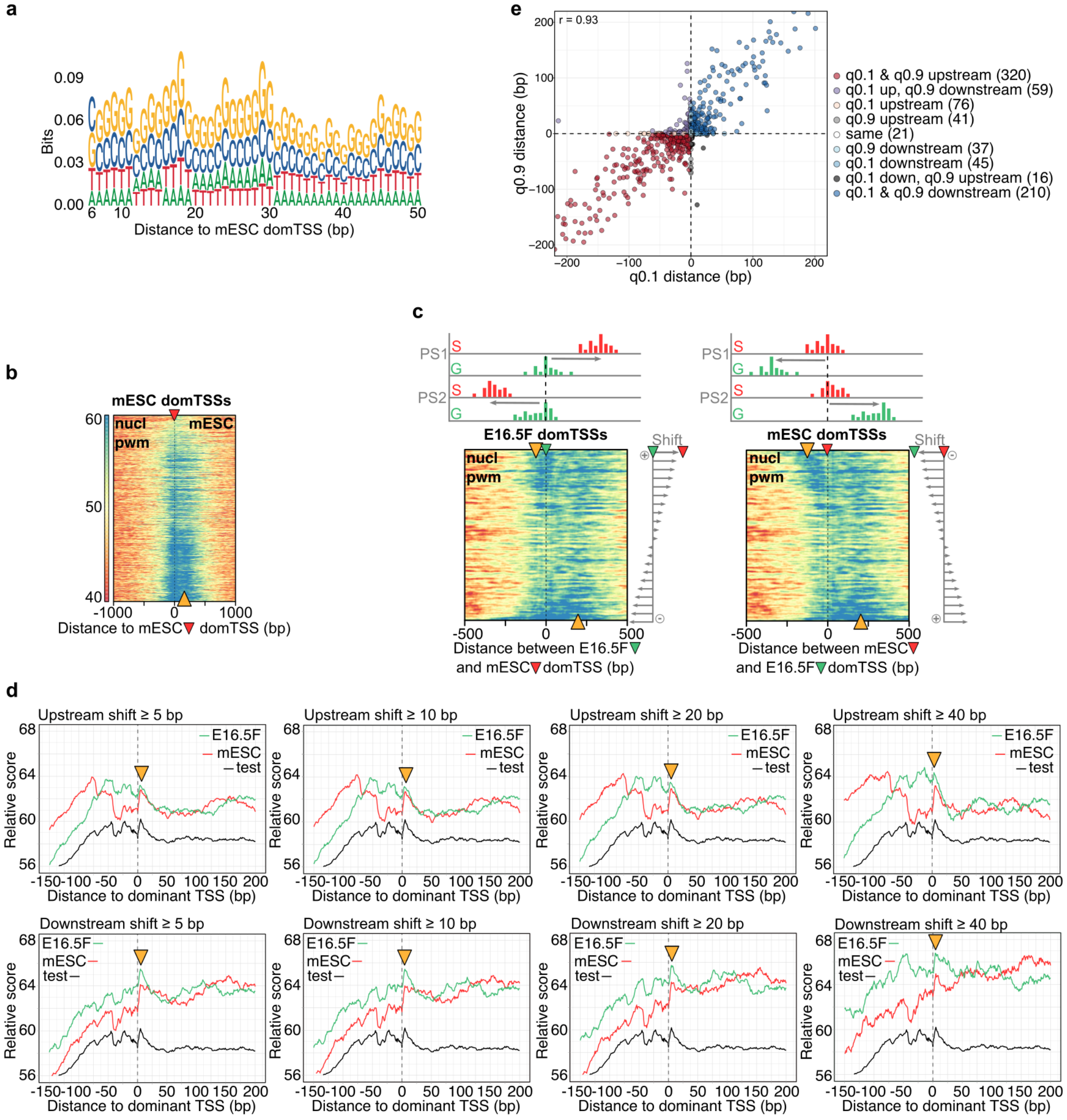


**Extended Data Figure 13. Sequence features underlying the precise +1 nucleosome positioning. a)** Sequence logo of a nucleosome preceding pwm constructed by aligning 8000 mESC broad promoter sequences centred on the dominant TSS. **b)** Heatmap visualisation of the nucleosome preceding pwm match scores in all mESC promoters centred on the dominant TSSs (marked with a red arrow on top) and ordered by the IQ-width (narrow promoters at the top, broad at the bottom of the heatmap). Scale reflects the percentile match score, windsorized to a 40-60 percentile match to more clearly depict the signal. **c)** Heatmap visualisation of the nucleosome preceding pwm matches in E16.5F vs mESC shifts centred on the E16.5F (left, marked with a green arrow) or mESC dominant CTSSs (right, marked with a red arrow) and ordered by the distance and orientation of the shift (schemes on top and right of the heatmaps). Scale is the same as in b). Orange arrowheads mark the high scoring areas. **d)** Metaplots of the nucleosome preceding pwm matches in: 1) E16.5F vs mESC shifts centred on the E16.5F (green) or mESC dominant CTSSs (red); 2) and mESC test broad promoters (excluded from pwm construction, black). Shifting promoters are divided into those with a dominant CTSS at least 5, 10, 20 or 40 bp upstream or downstream of the mESC dominant CTSSs. Orange arrowheads indicate the position of the nucleosome pwm match peak that aligns with the dominant TSSs. **e)** Scatter plot showing correlation of the distances between q0.1 or q0.9 tag cluster borders in E16.5F or mESC samples. q0.1 border marks the position of the $10^{th}$ percentile and the q0.9 border marks the position of the $90^{th}$ percentile of the tag cluster signal. $10^{th}$ and $90^{th}$ percentile signal positions are used to exclude outlier CTSSs and the effects of sequencing depth. High correlation of the q0.1 and q0.9 movements in shifting promoters (distances between two q0.1 or q0.9 borders in E16.5F compared to the mESC tag cluster) supports the +1 nucleosome repositioning as the underlying cause of the E16.5 shifts. The +1 nucleosome determines the "catchment area" within which transcription can initiate at multiple positions, with the dominant CTSS being at the optimal distance from the +1 nucleosome dyad. Therefore, repositioning of the +1 nucleosome is expected to influence both q0.1 and q0.9 borders in a coordinated manner - in the same direction and similar distance.

### Extended Data Figure 14.

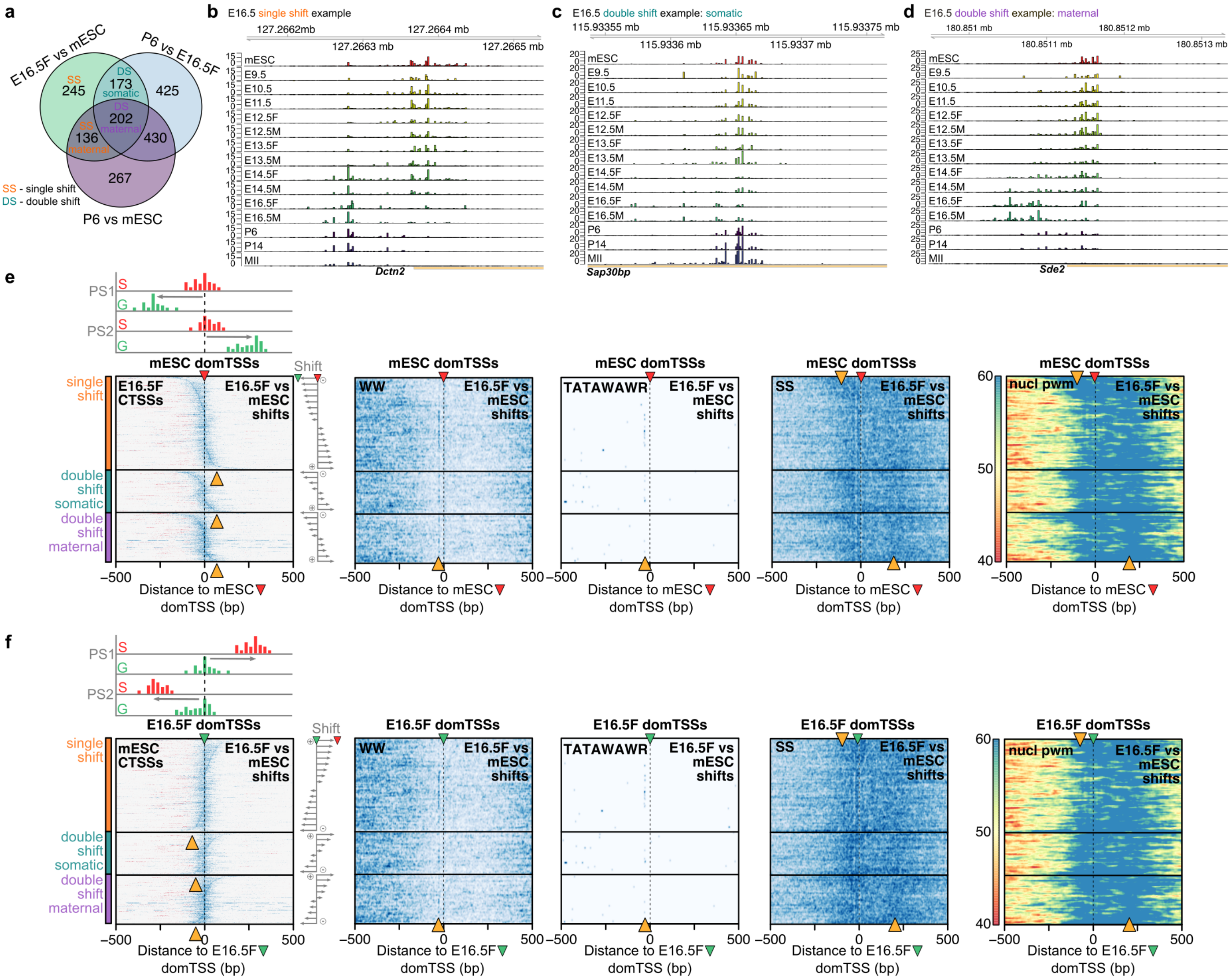

**Extended Data Figure 14. Types of shifts in late gonadal germ cells. a)** Venn diagram showing overlaps of shifting promoters identified between two named samples used to classify E16.5F vs mESC shifts into single and double shift promoters. Single shifts are identified as shifting between E16.5F and mESC, but not between E16.5F and the P6 oocyte. Double shifts are identified as shifting between E16.5F and mESC, and between E16.5F and P6 oocyte. Further, double shift somatic promoters exhibit CTSS patterns in the P6 oocyte highly similar to the mESC, i.e. they are not identified as shifting between the P6 oocyte and mESC. Double shift maternal promoters exhibit different CTSS patterns in all three stages - mESC, E16.5F and P6. **b-d)** CTSS patterns of example E16.5F promoters that exhibit a single or double shift - somatic or maternal. **e, f)** Sequence features of the E16.5F promoter shifting classes – heatmaps of CTSS coverage, WW, consensus TATA-box TATAWAWR motif, SS density or nucleosome-preceding pwm (generated in this study) match density. Promoters within each class are centred on the mESC dominant TSSs, marked with a red arrow e) or E16.5F dominant CTSS marked with a green arrow f) and ordered by the distance and orientation of the shift between E16.5F and mESC dominant CTSSs (schemes on top and on the right of the CTSS heatmaps). Orange arrowheads in CTSS heatmaps mark the inverted S shape of the CTSSs in each class, orange arrowheads in WW and TATAWAWR heatmaps indicate the expected position of the WW- or TATA-box (30 bp upstream of the dominant CTSS). Orange arrowheads in SS density heatmaps indicate positions of the SS enrichment, and the orange arrowheads in nucleosome pwm heatmaps indicate positions of the nucleosome-preceding pwm match density. None of the E16.5F shifting classes show a strong WW enrichment upstream of the of dominant CTSSs, hence transcription initiation in the E16.5F shifting promoter classes is not W-box dependent. However, the CTSS signal follows the SS enrichment and the nucleosome preceding pwm match density, indicating again that transcription initiation in E16.5F vs mESC shifting promoters is dictated by the +1 nucleosome positioning.

**Extended Data Figure 15.**

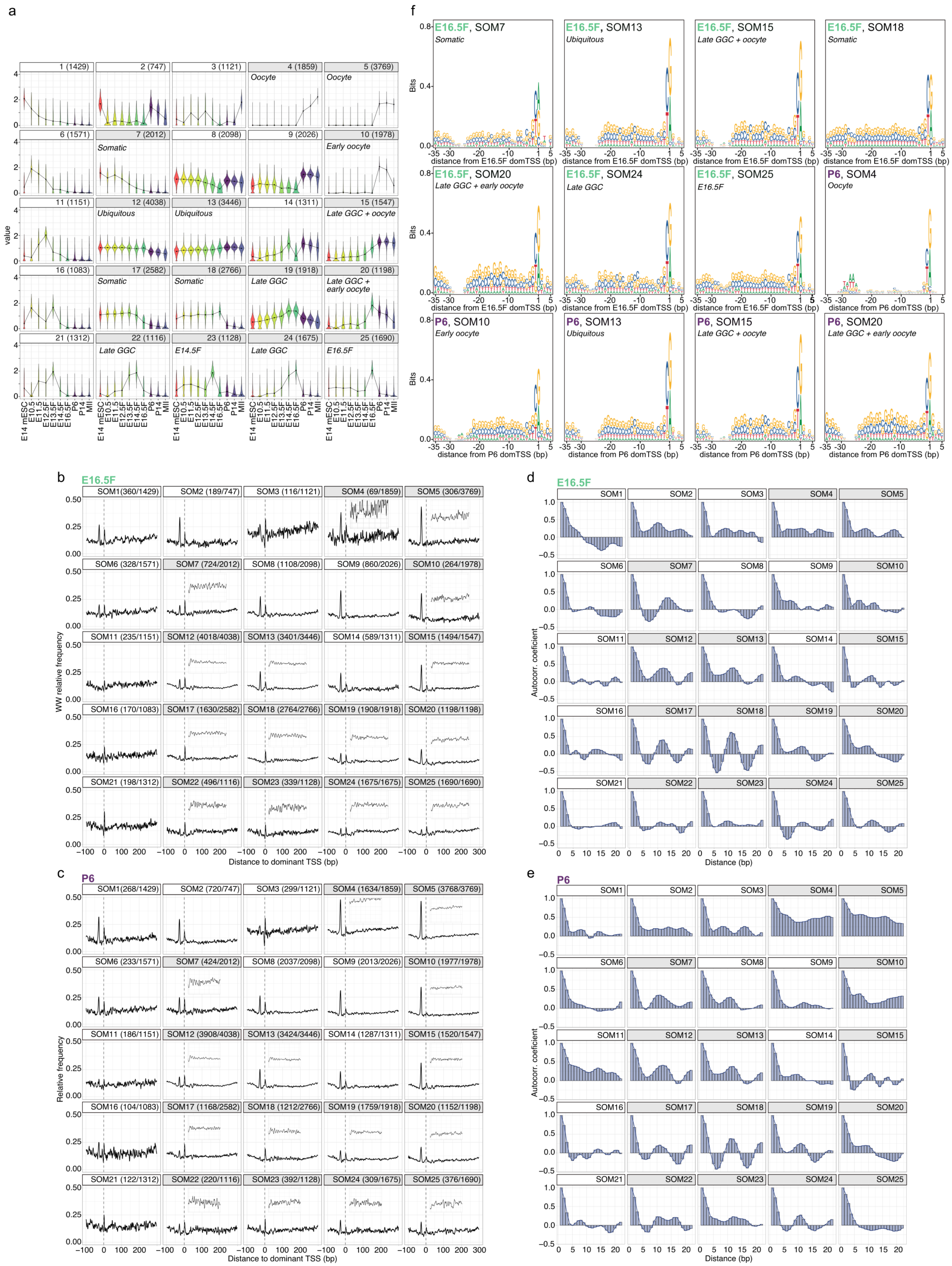
a
1 (1429)
2 (747)
3 (1121)
4 (1859)
Oocyte
5 (3769)
Oocyte
6 (1571)
7 (2012)
Somatic
8 (2098)
9 (2026)
10 (1978)
Early oocyte
11 (1151)
12 (4038)
Ubiquitous
13 (3446)
Ubiquitous
14 (1311)
15 (1547)
Late GGC + oocyte
16 (1083)
17 (2582)
Somatic
18 (2766)
Somatic
19 (1918)
Late GGC
20 (1198)
Late GGC + early oocyte
21 (1312)
22 (1116)
Late GGC
23 (1128)
E14.5F
24 (1675)
Late GGC
25 (1690)
E16.5F
value
E14 mESC
E10.5
E11.5
E12.5F
E13.5F
E14.5F
E16.5F
P6
P14
MII
f
E16.5F, SOM7
Somatic
E16.5F, SOM13
Ubiquitous
E16.5F, SOM15
Late GGC + oocyte
E16.5F, SOM18
Somatic
E16.5F, SOM20
Late GGC + early oocyte
E16.5F, SOM24
Late GGC
E16.5F, SOM25
E16.5F
P6, SOM4
Oocyte
P6, SOM10
Early oocyte
P6, SOM13
Ubiquitous
P6, SOM15
Late GGC + oocyte
P6, SOM20
Late GGC + early oocyte
Bits
distance from E16.5F domTSS (bp)
distance from P6 domTSS (bp)
b
E16.5F
SOM1(360/1429)
SOM2 (189/747)
SOM3 (116/1121)
SOM4 (69/1859)
SOM5 (306/3769)
SOM6 (328/1571)
SOM7 (724/2012)
SOM8 (1108/2098)
SOM9 (860/2026)
SOM10 (264/1978)
SOM11 (235/1151)
SOM12 (4018/4038)
SOM13 (3401/3446)
SOM14 (589/1311)
SOM15 (1494/1547)
SOM16 (170/1083)
SOM17 (1630/2582)
SOM18 (2764/2766)
SOM19 (1908/1918)
SOM20 (1198/1198)
SOM21 (198/1312)
SOM22 (496/1116)
SOM23 (339/1128)
SOM24 (1675/1675)
SOM25 (1690/1690)
WW relative frequency
Distance to dominant TSS (bp)
d
E16.5F
Autocorr. coeficient
Distance (bp)
c
P6
SOM1(268/1429)
SOM2 (720/747)
SOM3 (299/1121)
SOM4 (1634/1859)
SOM5 (3768/3769)
SOM6 (233/1571)
SOM7 (424/2012)
SOM8 (2037/2098)
SOM9 (2013/2026)
SOM10 (1977/1978)
SOM11 (186/1151)
SOM12 (3908/4038)
SOM13 (3424/3446)
SOM14 (1287/1311)
SOM15 (1520/1547)
SOM16 (104/1083)
SOM17 (1168/2582)
SOM18 (1212/2766)
SOM19 (1759/1918)
SOM20 (1152/1198)
SOM21 (122/1312)
SOM22 (220/1116)
SOM23 (392/1128)
SOM24 (309/1675)
SOM25 (376/1690)
Relative frequency
Distance to dominant TSS (bp)
e
P6
Autocorr. coeficient
Distance (bp)

**Extended Data Figure 15. Sequence features of the dominant CTSSs classified using SOM. a)** SOM-identified dominant CTSS clusters. Several example ubiquitous, somatic, oocyte-specific clusters are highlighted. The number of dominant CTSSs in each class is in brackets next to the SOM cluster number. **b)** WW relative frequency in E16.5F SOM clusters. The number of dominant CTSSs from SOM clusters expressed in the E16.5F stage is marked on the top of each SOM plot as expressed/total, where the total marks the total number of dominant CTSSs in that SOM cluster. The inset in the highlighted clusters shows a magnification of the 50-200 bp region. **c)** Same as b) albeit with dominant CTSSs from SOM clusters identified as expressed in the P6 sample. SOM clusters 7, 17, 18 (somatic, in b) E16.5F sample) and 12, 13 (ubiquitous, b) E16.5F or c) P6 sample) show strong WW-periodicity, in line with the dominant CTSS position being directed by the +1 nucleosome position. SOM clusters 19, 22 and 24 (late GGCs) and 25 (E16.5F) also display WW-periodicity, albeit weaker, presumably due to a lower number of sequences per cluster. On the contrary, clusters 4, 5 and 10 (oocyte specific, c) P6 sample) do not exhibit WW-periodicity, in line with its dominant CTSS position being dependent on a W-box (visible as a WW enrichment 30 bp upstream of the dominant CTSS position). **d)** Autocorrelation analysis of WW dinucleotide frequency in sequences 50-200 bp downstream of the dominant TSSs from SOM clusters expressed in the E16.5F sample. **e)** same as d) albeit using the dominant TSSs from SOM clusters expressed in the P6 oocyte. Autocorrelation analysis confirms the observations from b) and c) - clusters highlighted as exhibiting WW-periodicity, show the highest autocorrelation coefficient around 10-11 bp. **f)** Sequence logos encompassing 35 bp upstream and 5 bp downstream of the dominant TSSs (E16.5F or P6) in selected SOM clusters, as highlighted at the top of each plot. This sequence analysis confirms that oocyte specific clusters are W-box dependent (P6 SOM4 and 10).

### Extended Data Figure 16.

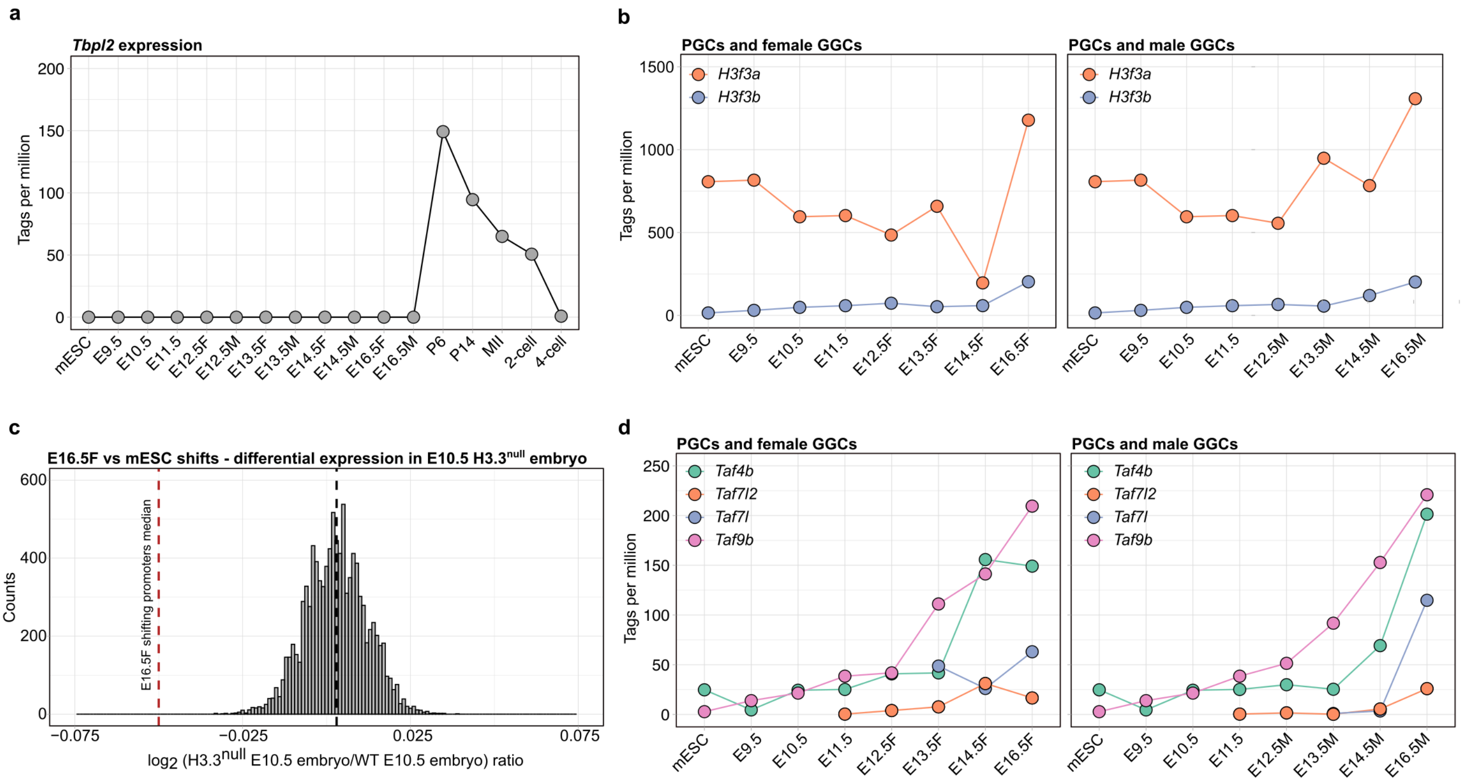


**Extended Data Figure 16. Histone H3.3 and alternative general transcription factor expression. a)** Expression of *Tbpl2* in mESC, PGCs, GGCs, oocyte and early embryos. **b)** Expression of two H3.3 genes *H3f3a* and *H3f3b* in mESC, primordial and gonadal female (left) and male (right) germ cells. Expression of both genes increases in male and female gonadal germ cells. **c)** Median log2-fold change of the E16.5F vs mESC shifting promoters (741, red dashed vertical line) that overlap with the genes differentially expressed in the E10.5M embryo (*Sox2*-cKO–*p53*$^{null}$ background) where the expression of the second exon of both H3.3 genes is completely abolished. The black dashed vertical line represents the median log2-fold change of 10000 iterations of 741 randomly selected genes from the differential expression data [2] (741 were chosen to match the size of the E16.5F shifting gene set). This data shows that the expression of E16.5F shifting promoters is influenced by H3.3. The effects of H3.3 removal on the transcriptome in the *p53*$^{null}$ background are limited, and only 5 % of transcribed genes in the dataset show statistically significant changes, therefore the whole set of differentially expressed genes was used. **d)** Expression of alternative GTFs *Taf4b*, *Taf7l2*, *Taf7l* and *Taf9b* in primordial and gonadal female (left) and male (right) germ cells. Their expression increases in both male and female gonadal germ cells.

**Extended Data Table 1.** Number of shifting TCs/promoters identified

| Sample 1 | Number of TCs | Shifting TCs[a] |
|---|---|---|
| mESC | 20863 | - |
| PGC E9.5 | 15886 | 678 |
| PGC E10.5 | 20425 | 239 |
| GGC E11.5 | 27018 | 279 |
| GGC E12.5F | 22953 | 292 |
| GGC E12.5M | 22577 | 239 |
| GGC E13.5F | 24265 | 337 |
| GGC E13.5M | 21932 | 351 |
| GGC E14.5F | 27420 | 481 |
| GGC E14.5M | 31655 | 404 |
| GGC E16.5F | 23087 | 829 |
| GGC E16.5M | 24431 | 707 |
| P6 | 35850 | 1098 |
| P14 | 33235 | 1130 |
| MII oocyte[b] | 35152 | 1028 |
| 2-cell embryo | 27335 | 1278 |
| 4-cell embryo[b] | 19705 | 1198 |

[a]Shifting TCs in each sample are identified in comparison with mESC.

[b]851 shifting promoters were identified in direct comparison of MII oocyte and 4-cell embryo.

**Extended Data References**


1. Veselovska, L. *et al.* Deep sequencing and de novo assembly of the mouse oocyte transcriptome define the contribution of transcription to the DNA methylation landscape. *Genome Biol* **16**, 209 (2015).
2. Jang, C.W., Shibata, Y., Starmer, J., Yee, D. & Magnuson, T. Histone H3.3 maintains genome integrity during mammalian development. *Genes Dev* **29**, 1377-92 (2015).
